# Polyamines sustain epithelial regeneration in aged intestines by modulating protein homeostasis

**DOI:** 10.1101/2024.07.26.605278

**Authors:** Alberto Minetti, Omid Omrani, Christiane Brenner, Gabriele Allies, Shinya Imada, Jonas Rösler, Saleh Khawaled, Feyza Cansiz, Sven W. Meckelmann, Nadja Gebert, Ivonne Heinze, Jing Lu, Katrin Spengler, Mahdi Rasa, Regine Heller, Omer Yilmaz, Alpaslan Tasdogan, Francesco Neri, Alessandro Ori

## Abstract

Aging hampers the regenerative potential of intestinal epithelium across species including humans, yet the underlying causes remain elusive. Here, using proteomic and metabolomic profiling of intestinal tissues together with functional assays, we characterized the temporal dynamics of regeneration following injury induced by 5-fluorouracil, a commonly used chemotherapeutic agent. Comparison of regeneration dynamics in mice of different ages revealed the emergence of a proteostasis stress signature and increased levels of polyamines following injury exclusively in old epithelia. Mechanistically, we show that delayed regeneration is an intrinsic feature of aged epithelial cells that display reduced protein synthesis and accumulation of ubiquitylated proteins. Notably, dietary restriction followed by re-feeding prior to injury increases polyamine pathway activation, enhances protein synthesis, and restores the regenerative capacity of aged intestines. Our findings highlight promising epithelial targets for interventions aimed at tackling the decline in tissue repair mechanisms associated with aging.

## Main text

The aging process involves a gradual decline of physiological integrity, resulting in diminished organ functionality, reduced regenerative capacities and increased risk of developing pathologies, including cancer ^1,2^. A long-standing question in aging biology and regenerative medicine is defining how tissue repair mechanisms change in the elderly, and which strategies can be pursued to ameliorate regeneration without heightening the risk of cancer onset. The small intestine is an excellent model to address these questions given its regenerative capacity mediated by tissue resident intestinal stem cells. Evidence from different groups has highlighted the contribution of cell-intrinsic alterations, changes in the communication with the stem cell niche, and systemic inflammation to the impaired function of old intestinal stem cells ^3–5^. However, although aging is known to delay the regenerative capacity of the small intestine following different types of injury ^6,7^, little is known about the temporal dynamics of the regeneration process and how these are affected in elderly individuals.

Maintenance of protein homeostasis, also known as proteostasis, is crucial for preserving cellular and tissue function over time, and its dysregulation has been proposed as a hallmark of aging ^8–10^. Mechanisms contributing to proteostasis impairment encompass, among others, alterations in protein turnover rates, a decline in protein degradation, and changes in protein quality control mechanisms ^1,11–13^. Notably, the requirements for proteostasis maintenance vary across different tissues and cell types ^12^. For instance, organs mainly composed of rarely dividing cells, e.g., the brain, are extremely prone to proteostasis impairment and tend to accumulate protein aggregates with age ^14,15^. Conversely, in proliferative tissues such as the intestinal epithelium, the evidence for protein homeostasis impairment during aging is less described, and it mainly arises following stress, e.g., upon tissue damage ^16,17^.

Polyamines are small by-products of food metabolism, and they are gaining relevance in the context of protein homeostasis maintenance and aging. Polyamines are known to orchestrate important aspects of proteostasis including translation elongation and termination, and autophagy ^18–21^. Moreover, recent evidence shows polyamines controlling translation rates of hair follicle stem cells, where changes in protein synthesis rates influence stem cell fates ^22^, highlighting their importance in regulating stem cell adaptation. Spermidine, one of the most active polyamines, enhances translation efficiency via hypusination of the eukaryotic initiation translation factor 5A (EIF5A) ^23^. Hypusinated EIF5A has been shown to both alleviate stalling on motifs containing poly-proline tracts and stimulate peptidyl-tRNA hydrolysis in translation termination, thus improving translation efficiency ^18–21^. Polyamines are also reported to tightly regulate other biological processes, including cellular proliferation, mitochondrial respiration, and immune responses ^24,25^. Moreover, despite differences among tissues, their abundance seems to decrease with age ^26^, thus raising attention to their relevance for healthy aging and cytoprotection. Strategies aimed to improve intestinal regeneration have been proposed including dietary interventions based on dietary restriction (DR) and fasting ^4,6,7,27–30^. Interestingly, in the context of dietary interventions, recent findings showed that post-fast re-feeding improves intestinal stem cell-mediated regeneration via polyamine metabolism ^31^. Whether these pathways contribute to age-related differences in intestinal epithelium regeneration remains elusive.

To model small intestine regeneration, we adopted 5-Fluorouracil (5-FU), a widely used chemotherapeutic agent known to inhibit DNA replication, RNA synthesis, and affect ribosome function ^32–35^. Within the gastrointestinal system, 5-FU induces intestinal mucositis, leading to decreased body weight primarily due to fluid imbalance and food malabsorption. We chose 5-FU because of (i) the mechanism of action targeting both nucleic acids and protein synthesis machinery, (ii) the relatively milder effects compared to alternative injury models like DSS or γ-irradiation, and (iii) the established age-related differences in intestinal regeneration ^6^. Here, investigating the dynamics of intestinal regeneration in young and old mice following a single injection of 5-FU, we found that upon injury old intestinal epithelia promote polyamine metabolism to overcome the emergence of proteostasis stress induced by tissue damage. Additionally, we show that the regenerative capacity of old intestines can be restored by activating the polyamine pathway before injury via an intervention based on dietary restriction followed by re-feeding.

### Proteome dynamics during intestinal epithelium regeneration

In order to dissect the temporal dynamics of small intestine regeneration, we collected whole tissue samples from small intestines of male mice on days 2, 5 and 7 after a single injection of 5-FU or vehicle control (PBS) (Fig. 1a). In young mice (3-4 months), 5-FU induced a mild reduction in body weight (∼5%) that peaked at 2 days post-injection (dpi) and resolved at 5 dpi (Fig. 1b). The effect on body weight was accompanied by histological changes of the intestinal epithelium including a decrease in crypt number (Fig. 1c, d), density of differentiated cells within the villi (Fig. 1c, e), number of phospho histone H3 (pH3)-positive cells (Fig. 1f), and crypt length (Fig. S1a). Although slight variations in temporal dynamics were observed among the different histological parameters, they were all consistent with reduced cell proliferation and tissue renewal at 2 dpi that typically resolved between 5 and 7 dpi.

**Figure 1.**
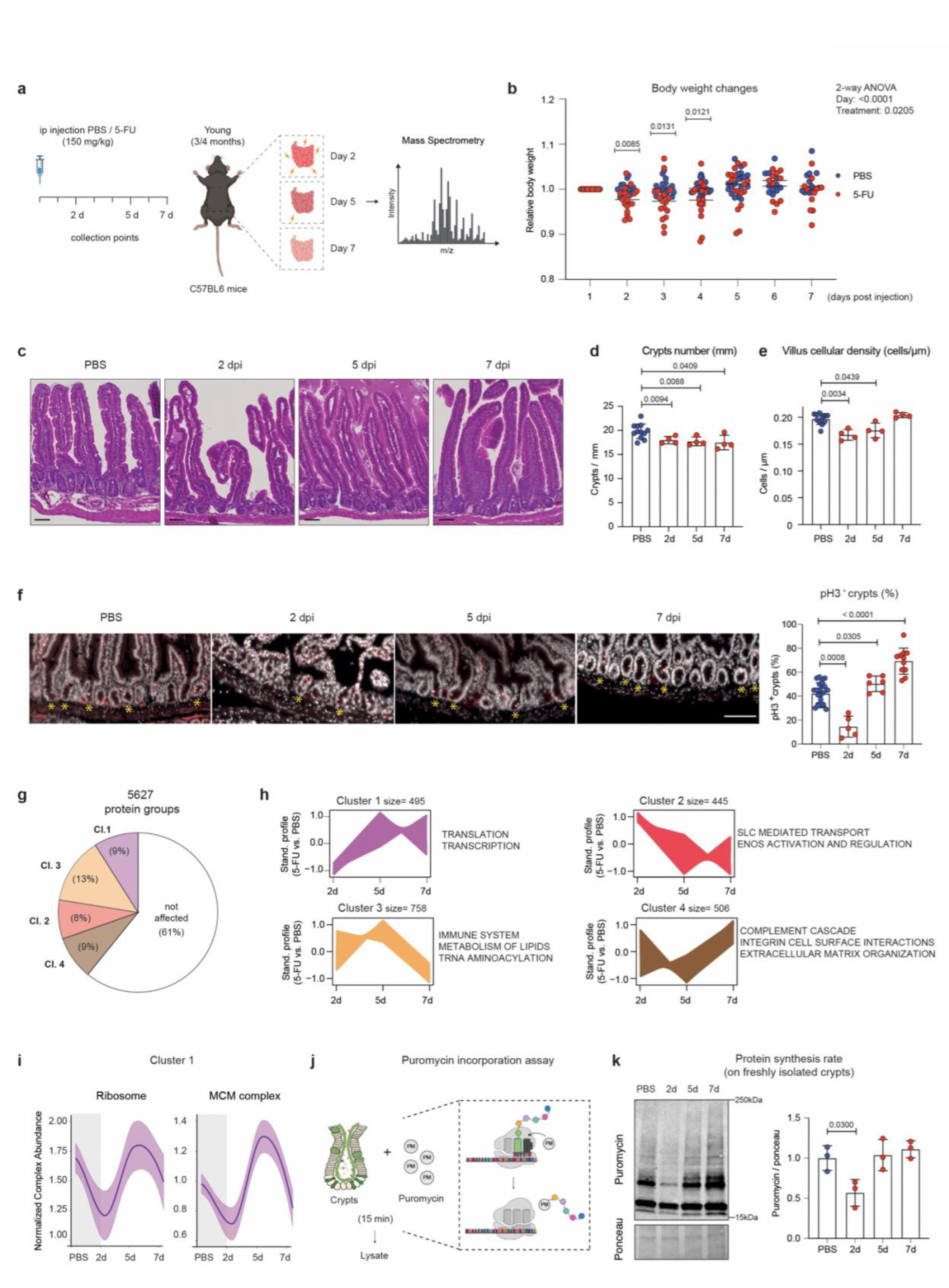
Identifying proteome dynamics during regeneration of young small intestines. **a** Schematic of 5-FU induced regeneration. ip: Intraperitoneal. **b** Relative body weight of young mice treated with a single dose of 5-FU or PBS as control. Body weight of each mouse was normalized to its body weight at the time of injection. *n*=18 PBS, *n*=17 5-FU mice. Black line indicates the median for each day in each treatment group. *P*-value is calculated with Welch’s t test by comparing the average body weight of 5-FU treated with the PBS-treated mice on the indicated day, and by two-way ANOVA for overall day and treatment comparisons. **c** Representative pictures of hematoxylin and eosin (H&E) staining of the small intestine from indicated treatments and time points. Scale bar, 50 µm. **d** Quantification of number of crypts per millimeter of small intestine in the indicated groups. PBS treated mice from different days were combined. Each dot represents one mouse. *n*=12 PBS, *n*=4 5-FU mice per group. Error bars represent the SD. *P*-value was calculated by Welch’s t-test. **e** Quantification of number of cells per one micrometer of villi in the indicated groups. PBS-treated mice from different days were combined. Each dot represents one mouse. *n*=11 PBS, *n*=4 5-FU mice per group. Error bars represent the SD. *P*-value was calculated by Welch’s t-test. **f** Left: Representative pictures of pH3 staining from indicated treatment and time point. Scale bar, 100 µm. Asterisks indicate the pH3+ crypts. Right: Percentage of pH3+ crypts in the indicated groups. PBS-treated mice from different days were combined. Each dot represents one mouse. *n*=5-24 mice per group. Error bars represent the SD. *-* value was calculated by Welch’s t-test. **g** Classification of protein groups quantified by proteomics according to their abundance changes relative to PBS controls. **h** Abundance profiles for the 4 clusters of proteins affected by 5-FU. Representative REACTOME gene sets significantly enriched in each cluster are shown. **i** Abundance profile of ribosome (67 proteins) and MCM (6 proteins) complexes. The protein abundances of individual complex members were normalized to the median complex abundance of PBS-treated mice, and profiles plotted using a loess smooth function. The shaded area around the regression line represents the 95% confidence interval. **j** Scheme of puromycin incorporation assay performed on freshly isolated crypts. **k** Left: Representative immunoblot for puromycin incorporation. Right: Quantification of puromycin incorporation into proteins relative to ponceau staining (loading control). Each dot represents one mouse. *n*=3 mice per group. Error bars represent the SD. *P*-value was calculated by Welch’s t-test.

**Figure S1.**
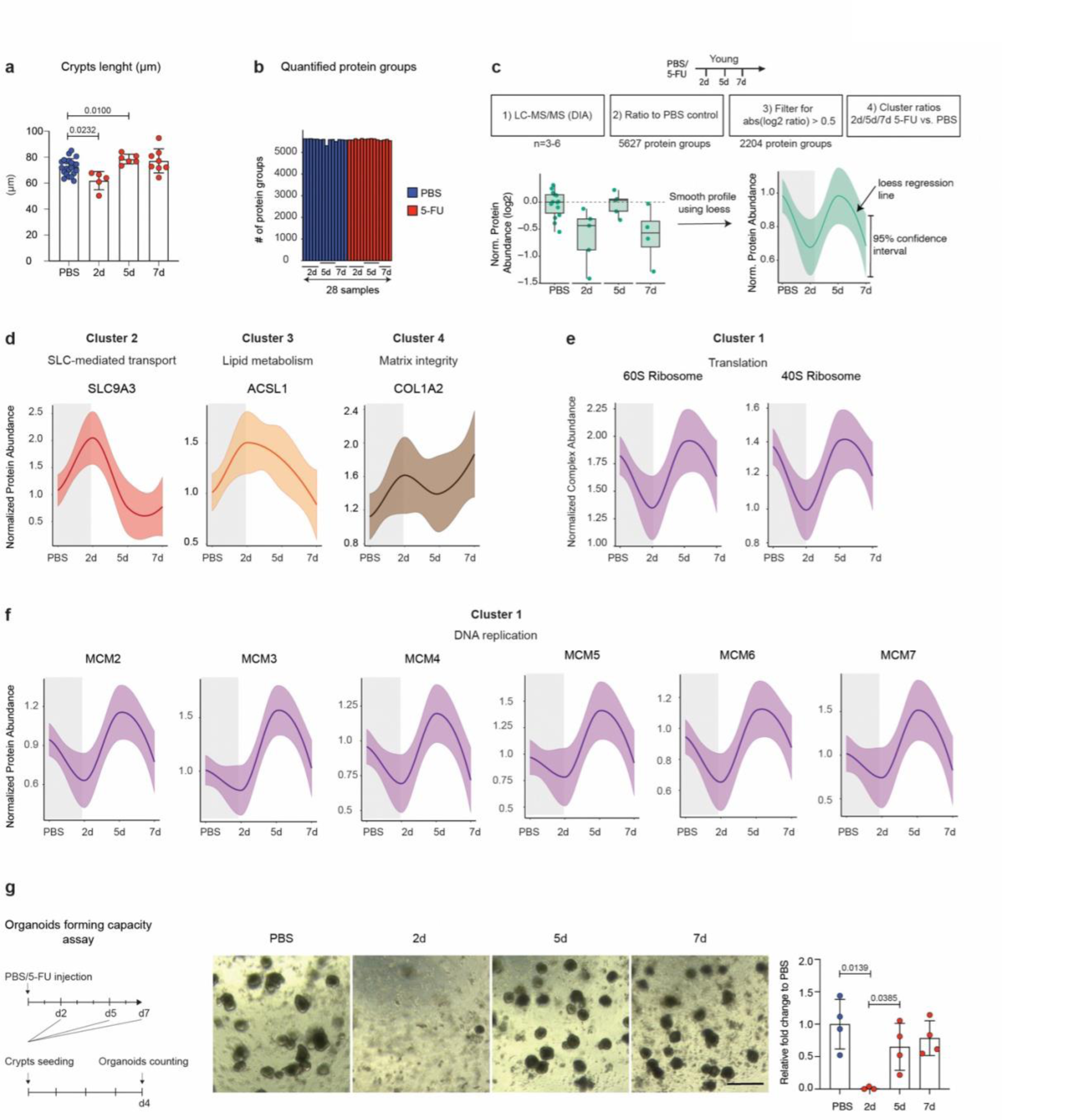
Proteome dynamics during young small intestinal epithelium regeneration. **a** Average length of the crypts from young mice in the indicated groups. PBS-treated mice from different days were combined. Each dot represents one mouse. *n*=5-22 mice per group. Error bars represent the SD. *P* value was calculated by Welch’s t test. **b** Number of quantified protein groups in the indicated treatments and time points. **c** Schematic of experimental design, proteomics data analysis and visualization strategy. **d** Representative abundance profiles for proteins affected by 5-FU treatment and belonging to different clusters. **e** Protein abundance profiles of 40S (26 proteins) and 60S (41 proteins) ribosome complexes. **f** Protein abundance profiles of components of the MCM complex. **g** Left: Schematic of colony forming assay. Middle: quantification of the growing organoids in the indicated groups normalized to the average of PBS, scale bar 10 µm. *n*=3/4 mice per group. Error bars represent the SD. *P*-value was calculated by Welch’s t-test.

To monitor the dynamics of intestinal regeneration at the proteome level, we analyzed samples from the whole intestinal tissue using label-free Data Independent Acquisition (DIA) mass spectrometry. We quantified 5627 protein groups with at least 2 proteotypic peptides across all the time points and replicates (Fig. S1b). We then used k-means clustering to group proteins that show changes in abundance following 5-FU injection relative to vehicle control (Fig. S1c). The majority (>60%) of protein groups quantified showed no change in their abundance. We found 2204 proteins affected by 5-FU that could be classified into 4 major clusters according to their abundance profile (Fig. 1g) (Table S1). The 4 clusters were enriched for specific pathways (Fig. 1h). These included pathways related to transcription and translation, lipid metabolism, immune response and extracellular matrix organization, among others (Fig. 1i, S1d). Many of these pathways have been functionally linked to intestine regeneration ^36–42^.

To functionally validate our findings, we focused on cluster 1, which includes multiple components of the minichromosome maintenance (MCM) complex and the ribosome (Fig. 1i, S1e,f). We confirmed reduced protein synthesis on freshly isolated intestinal crypts at 2 dpi using *ex-vivo* puromycin incorporation assay (Fig. 1j, k). In line with reduced protein synthesis, crypts from 2 dpi displayed a drastically reduced organoid-forming capacity that was completely restored at 5 dpi (Fig. S1g). Together, these data define histological and molecular signatures that characterize different phases of intestinal regeneration and highlight a temporal regulation of the proteome following injury by 5-FU.

### Emergence of proteostasis stress in old intestines upon injury

To investigate how aging affects proteome dynamics during intestinal epithelium regeneration, we performed the same injury experiment using old (22-26 months) mice. Consistent with previous studies ^4,6,7,27,28,30^, we found that aged mice displayed more pronounced body weight loss (∼10% vs. 5% in the young) and required 7 days to restore their body weight to a level comparable to the ones of vehicle-treated controls (Fig. 2a and S2a). At the histological level, the old epithelia exhibited different dynamics for crypts number (Fig. 2b), number of pH3-positive cells (Fig. 2c), density of differentiated cells within the villi (Fig. S2b), but not crypt length (Fig. S2c), compared to the young counterparts. In order to gain insights into the molecular mechanisms of delayed regeneration, we acquired proteomics data from the whole intestinal tissue at the same time points post 5-FU injection collected for young animals. For each age group and time point, we calculated fold changes relative to vehicle controls and compared protein abundance changes between young and old mice using multivariate analysis of variance (MANOVA) (Fig. 2d). Using k-means clustering, we could identify four major clusters of proteins that displayed different abundance changes in old vs young (446 proteins in total, Fig. 2e) (Table S2). Pathway enrichment analysis revealed a network of proteins related to protein homeostasis (hereafter proteostasis) that displayed decreased levels at 2 dpi and increased levels at 7 dpi in old mice compared to young ones (Fig. 2e, f). This cluster included multiple members of the 19S regulatory particle of the proteasome and of the chaperonin-containing TCP-1 complex (CCT), translation initiation factors (EIF1B, EIF2B1, EIF4A2), and autophagy-related proteins including ATG5 and ATG7 (Fig. 2f, g). Other proteins related to proteostasis also showed significantly different dynamics in the old vs. young intestine, although with different profiles (Fig. S2d). Interestingly, other proteostasis network components such as ribosomes (Fig. 2g) and 20S proteasome (Fig. S2e), as well as the MCM complex (Fig. S2f), showed identical dynamics in both young and old mice. We confirmed some of these findings by immunoblot against the autophagy receptor p62 (SQSTM1) and the ribosomal protein RPS6 in an independent set of freshly isolated intestinal crypts (Fig. S2g, S2h, S2i). We additionally confirmed that p62 protein level changes were independent of mRNA expression (Fig. S2j). In agreement with an impairment of proteostasis, we found increased levels of proteins modified by polyubiquitin chains linked via lysine 48 (K48) at 5 dpi in old mice (Fig. 2i), a marker for protein degradation, but not total ubiquitin (Fig. S2k). Conversely, *ex-vivo* measurements of protein synthesis were overall similar in young and old crypts (Fig. 2h), consistent with the similar dynamics of ribosomal proteins (Fig. 2g). Together, our data confirm delayed intestinal regeneration in old mice following 5-FU injury. In addition, we found that delayed regeneration is accompanied by the emergence of a proteostasis stress signature that affects the temporal dynamic of specific proteins in the aged intestinal epithelium and crypts.

**Figure 2.**
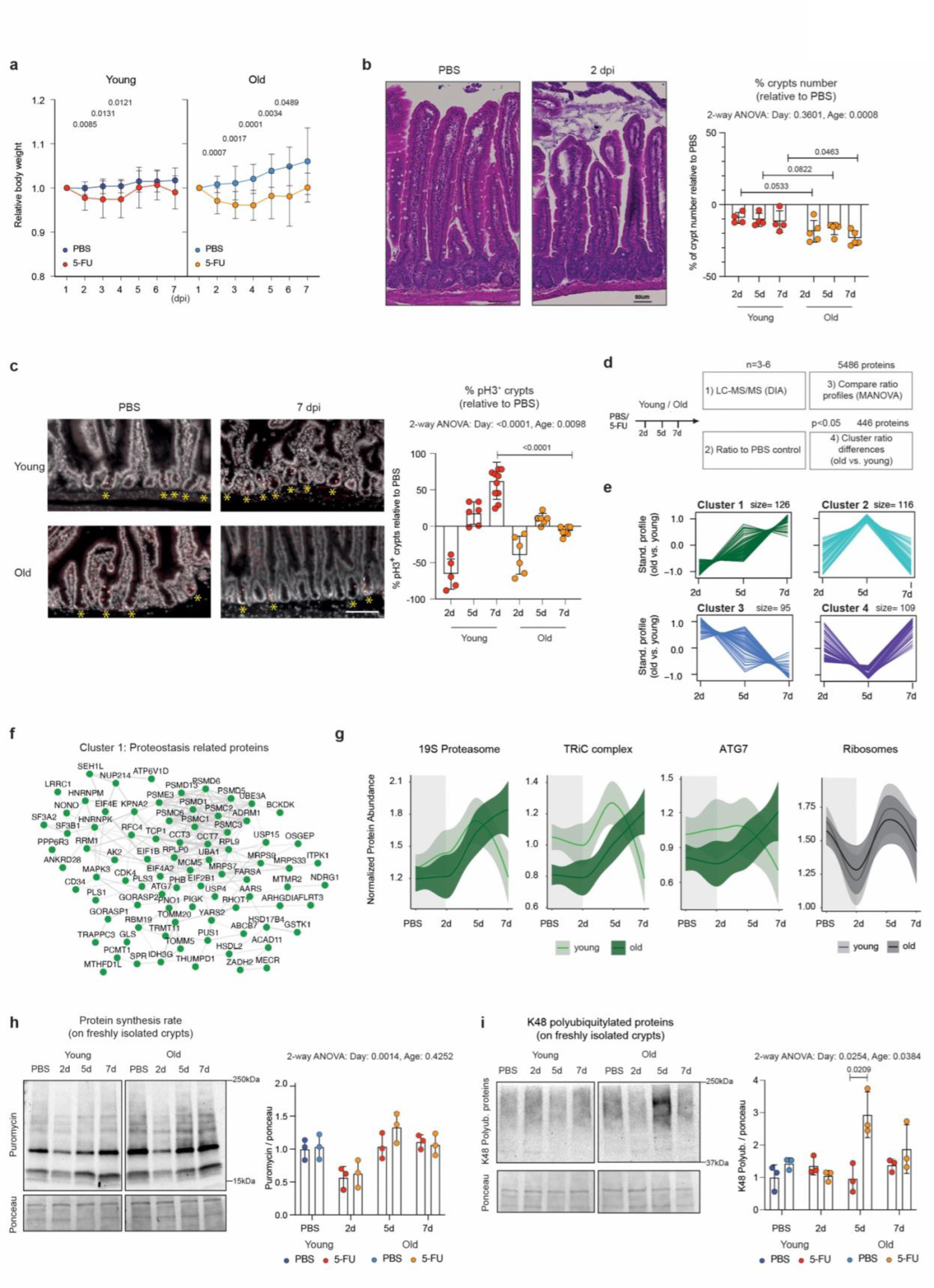
Old mice show signs of proteostasis stress in the intestinal tissue upon injury. **a** Relative body weight of young (left) and old (right) mice treated with a single dose of 5-FU or PBS as control. Body weight of each mouse was normalized to its body weight at the time of injection. *n*=18 PBS, *n*=17 5-FU young mice and *n*=12 PBS, *n*=14 5-FU old mice. *P*-value is calculated with Welch’s t-test by comparing the average body weight of 5-FU treated with the PBS treated mice on the indicated day. **b** Left: Representative pictures of H&E staining of small intestine. Scale bar, 50 µm. Right: quantification of number of crypts per millimeter of small intestine expressed as percentage relative to average PBS control mice. *n*=5/15 mice per group. Error bars represent the SD. *P*-value was calculated using Welch’s t-test for time point comparison and two-way ANOVA for overall day and age comparisons. **c** Left: Representative pictures of pH3 staining from indicated treatments and time points. Scale bar, 100 µm. Asterisks indicate the pH3+ crypts. Right: Change in number of pH3^+^ crypts expressed as percentage relative to average PBS control mice. *n*=5-11 mice per group. Error bars represent the SD. *P*-value was calculated using Welch’s t-test for time point comparison and two-way ANOVA for overall day and age comparisons. **d** Workflow for the comparison of protein abundance profiles between young and old mice. **e** Fold change profiles (old vs. young, log2) for the 4 clusters of proteins that display different dynamics in young and old mice upon 5-FU treatment. **f** Network of proteostasis related proteins from cluster 1. Protein-protein interactions were derived from STRING ^43^ using a cut-off of 0.7. **g** Protein abundance profiles for components of the 19S proteasome (24 proteins), TriC (7 proteins) complexes, ATG7 and ribosomal proteins (67 proteins). **h** Left: Representative immunoblot for the puromycin incorporation. Right: Quantification of puromycin incorporation. Each dot represents one mouse. *n*=3 mice per group. Error bars represent the SD. *P*-value was calculated using Welch’s t-test for time point comparison and two-way ANOVA for overall day and age comparisons. **i** Left: Representative immunoblot for the K48 polyubiquitylated proteins. Right: Quantification of K48 polyubiquitylated proteins relatively to ponceau staining (loading control). Each dot represents one mouse. *n*=3 mice per group. Error bars represent the SD. *P*-value was calculated using Welch’s t-test for time point comparison and two-way ANOVA for overall day and age comparisons.

**Figure S2.**
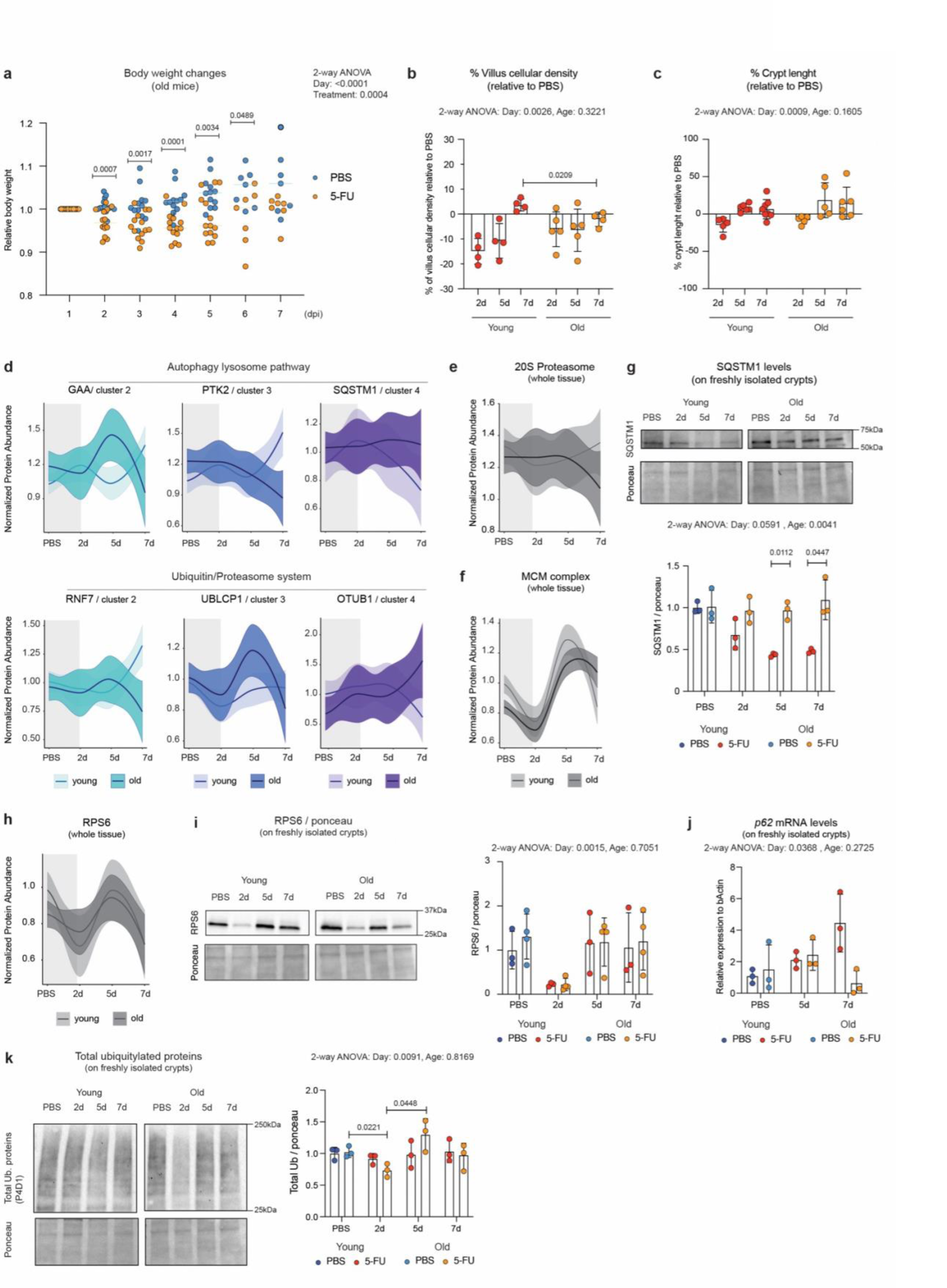
Perturbation of proteostasis after injury in aged intestinal tissue. **a** Relative body weight of old mice treated with a single dose of 5-FU or PBS as control. Body weight of each mouse was normalized to its body weight at the day of injection. *n*=12 PBS, *n*=14 5-FU mice. Black line indicates the median for each day in each treatment group. *P*-value is calculated with Welch’s t-test by comparing the average body weight of 5-FU treated with the PBS-treated mice on the indicated day, and by two-way ANOVA for overall day and treatment comparisons. **b** Percentage of villus cellular density relative to PBS. In each time point 5-FU values were normalized to the average of PBS and expressed as percentage. Each dot represents one mouse. *n*=4-11 mice per group. Error bars represent the SD. *P*-value was calculated using Welch’s t-test for time point comparison and two-way ANOVA for overall day and age comparisons. **c** Percentage of crypts length relative to PBS. In each time point 5-FU values were normalized to the average of PBS and expressed as percentage. Each dot represents one mouse. *n*=5-22 mice per group. Error bars represent the SD. *P*-value was calculated using two-way ANOVA for overall day and age comparisons. **d** Upper: Protein abundance profiles of the proteins related to the autophagy lysosome pathway. Lower: Protein abundance profiles of the ubiquitin proteasome pathway. Proteostasis-related proteins list was obtained from https://www.proteostasisconsortium.com/pn-annotation/ **e-f** Protein abundance profiles of the 20S proteasome (17 proteins) and MCM complex (6 proteins). **g** Upper: representative immunoblot for the SQSTM1 protein in the indicated groups. Lower: Quantification of SQSTM1 protein level relative to ponceau staining. Each dot represents one mouse. *n*=3 mice per group. Error bars represent the SD. *P*-value was calculated using Welch’s t-test for time point comparison and two-way ANOVA for overall day and age comparisons. **h** Protein abundance profiles of the RPS6 protein. **i** Left: representative immunoblot for the RPS6 protein. Right: Quantification of RPS6 protein level in the indicated groups. Each dot represents one mouse. *n*=3-4 mice per group. Error bars represent the SD. *P*-value was calculated using Welch’s t-test for time point comparison and two-way ANOVA for overall day and age comparisons. **j** Relative mRNA expression level of the *p62* gene in the indicated groups. Error bars represent the SD. *P*-value was calculated using Welch’s t-test for time point comparison and two-way ANOVA for overall day and age comparisons. **k** Left: representative immunoblot for the total ubiquitylated conjugates. Right: Quantification of total ubiquitylated proteins relative to ponceau staining in the indicated groups. Each dot represents one mouse. *n*=3 mice per group. Error bars represent the SD. *P* value was calculated by Welch’s t test. *P*-value was calculated using Welch’s t-test for time point comparison and two-way ANOVA for overall day and age comparisons.

### Delayed regeneration of old intestinal organoids

Both cell-intrinsic and extrinsic factors influence intestinal regeneration ^4,6,7^. To assess to what extent the delayed regeneration of old intestines and the associated proteostasis stress have a cell-intrinsic component, we used intestinal organoids to model the 5-FU injury in culture. Small intestinal organoids from young and old mice were treated with different concentrations of 5 -FU or vehicle control for 24 h, passed to wash out 5-FU, and their morphological composition was assessed 3 days later (Fig. 3a). First, we showed that, as expected, 5-FU treatment induced dose-dependent changes in the proportion of cystic, budded, or apoptotic organoids both in young and old (Fig 3b, c). However, old organoids displayed greater sensitivity to 5-FU. At the lowest concentration tested (2.5 and 1.7 µg/ml of 5-FU), young organoids were able to generate cystic/budded/apoptotic organoids in a comparable proportion to their PBS control, whereas old organoids showed a significant decrease of budded organoids already at these drug concentrations (Fig. 3b, c). To further define whether intestinal organoids recapitulated other signatures of delayed regeneration observed *in vivo*, we selected a specific concentration of 5-FU (2.5 µg/ml), which was the highest concentration displaying age-related morphological differences, and then tested by immunoblot some of the key parameters emerging from *in vivo* analysis. We first checked apoptosis induction to evaluate whether the extent of damage induced by 5-FU was different between young and old organoids. Cleaved Caspase 3 (cCasp3) to Caspase 3 (Casp3) ratio was similarly elevated 3 days after 5-FU wash out in organoids from young and old mice (Fig. 3d), indicating a comparable induction of apoptosis. Next, we tested organoids proliferation by pH3 immunoblot. As expected, the levels of pH3 were drastically reduced 24 h after 5-FU treatment in both young and old organoids. However, young organoids recovered pH3 levels on day 3 after 5-FU wash out, while old organoids did not (Fig. 3e). Consistent with reduced proliferation 3 days post injury but differently from freshly isolated crypts (Fig. S2g), old organoids also displayed reduced protein synthesis levels compared to young, as assessed by puromycin incorporation (Fig. 3f). Importantly, the delayed regeneration of old organoids post 5-FU injury was accompanied by some of the proteostasis stress signatures that we observed *in vivo,* including increased levels of p62/SQSTM1 and K48-polyubiquitylated proteins (Fig. 3g, h). Together, these data show that old intestinal organoids possess reduced regenerative capacity following 5-FU treatment and show signs of proteostasis stress, suggesting an epithelial cell-intrinsic component to the delayed intestinal regeneration observed *in vivo*.

**Figure 3.**
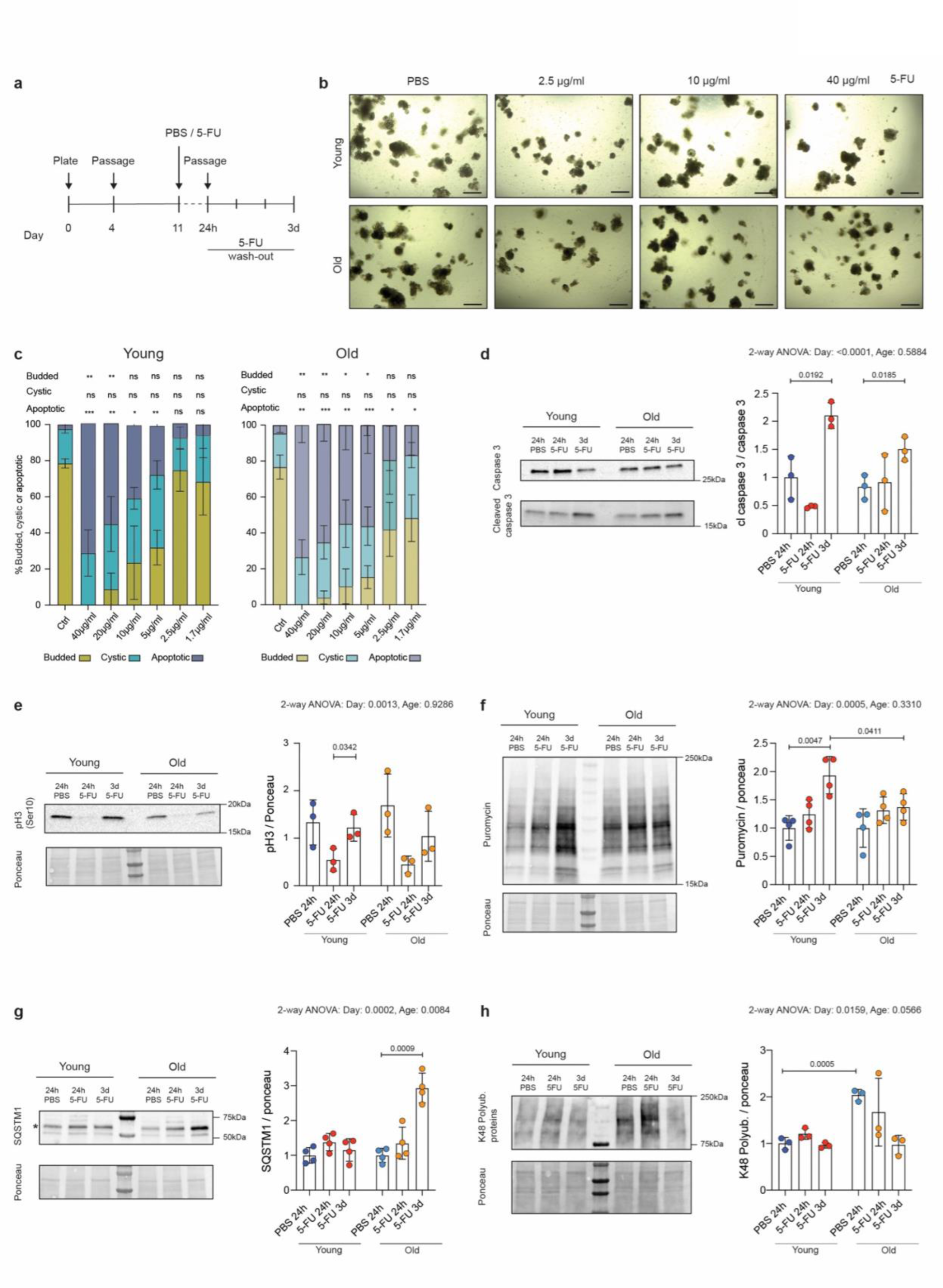
Delayed regeneration of organoids from old mice upon treatment with 5-FU. **a** Scheme of the 5-FU-induced regeneration experiment using intestinal organoids. **b** Brightfield images of intestinal organoids in the indicated groups. Pictures were taken 3 days after PBS or 5-FU wash out. Scale bar 200 µm. **c** Quantification of organoid morphology at day 3 post 5-FU treatment. *n*=3 mice per group. Error bars represent the SD. *P*-value was calculated by Welch’s t-test comparing the different conditions to young or old PBS control. * = *p* < 0.05, ** = *p* < 0.01, *** = *p* < 0.001. **d** Left: representative immunoblot for cleaved and total caspase 3. Right: quantification of caspase 3 immunoblot in the indicated groups. Each dot represents one mouse. *n*=3 mice per each indicated condition. Error bars represent the SD. *P*-value was calculated using Welch’s t-test for time point comparison and two-way ANOVA for overall day and age comparisons. **e** Left: representative immunoblot for the pH3 protein. Right: Quantification of the pH3 level normalized to ponceau in the indicated groups. Each dot represents one mouse. *n*=3 mice per each indicated condition. Error bars represent the SD. *P*-value was calculated using Welch’s t-test for time point comparison and two-way ANOVA for overall day and age comparisons. **f** Left: representative immunoblot for the puromycin incorporated proteins. Right: Quantification of the puromycin incorporated proteins in the indicated groups. Each dot represents one mouse. *n*=4 mice per each indicated condition. Error bars represent the SD. *P*-value was calculated using Welch’s t-test for time point comparison and two-way ANOVA for overall day and age comparisons. **g** Left: representative immunoblot for the SQSTM1 protein. Right: quantification of SQSTM1 protein level normalized to ponceau in the indicated groups. Each dot represents one mouse. *n*=4 mice per each indicated condition. Error bars represent the SD. *P*-value was calculated using Welch’s t-test for time point comparison and two-way ANOVA for overall day and age comparisons. **h** Left: representative immunoblot for the K48 polyubiquitylated proteins. Right: quantification of K48 polyubiquitylated proteins in the indicated groups. Each dot represents one mouse. *n*=3 mice per condition. Error bars represent the SD. *P*-value was calculated using Welch’s t-test for time point comparison and two-way ANOVA for overall day and age comparisons.

### The polyamine pathway is induced in aged intestinal epithelia following damage

Recent data highlighted that fasting and re-feeding modulate protein synthesis capacity in the intestinal epithelium of young mice due to the mechanistic target of rapamycin complex 1 (mTORC1)-driven regulation of the polyamines pathway ^31^. Intrigued by these findings and given the influence of polyamines on tissue regeneration ^24,44^, we asked whether similar mechanisms might contribute to influencing intestinal regeneration in old mice (Fig. 4a). Thus, we investigated the dynamics of key enzymes involved in polyamine metabolism in our proteome datasets (Fig. 4b). We found ornithine aminotransferase (OAT) to be reduced 2 dpi in both young and old tissues (Fig. 4c), concomitantly with reduced ribosomes and protein synthesis. Interestingly, carbamoyl phosphate synthetase 1 (CPS1) and ornithine transcarbamoylase (OTC), the two enzymes involved in the first steps of the urea cycle, were reduced 5 dpi in the old (Fig. S3a). Conversely, polyamine oxidase (PAOX), deoxyhypusine synthase (DHPS) and deoxyhypusine hydroxylase (DOHH) showed an increase of abundance at 5 dpi that was more pronounced in the old tissues (Fig. 4c). These data suggest an activation of polyamine metabolism over urea cycle in aged epithelia following injury, a pattern that closely resembles the observations made by Imada *et al.* following re-feeding. Therefore, we decided to adopt the same targeted metabolomic approach used by Imada *et al.* to measure polyamines and related metabolites in freshly isolated crypts. We found comparable levels of polyamines in young and old uninjured crypt samples (Fig. 4d). However, polyamines levels, differently from urea cycle metabolites (Fig. S3b), varied considerably between young and old following 5-FU injection, with a characteristic increase in abundance at 5 dpi exclusively in aged mice (Fig. 4c) that coincided with the induction of key enzymes of the pathway (Fig. 4b). To assess the functional relevance of these findings, we evaluated the total and hypusinated levels of EIF5A, the main downstream effector of the polyamine pathway. The levels of both followed a dynamic consistent with the observed levels of polyamines being increased at 5 dpi in old mice (Fig. 4e, S3c). Given the known role of hypusinated EIF5A in alleviating ribosome stalling at specific motifs including polyproline tracts ^20^, we decided to investigate further the dynamics of proteins enriched in these motifs (Fig S3d). We found several proteins containing these motifs to show distinct dynamics in young and old intestines (Fig. 4f). In particular, proteins containing multiple EIF5A-dependent motifs were over-represented in cluster 2 of our MANOVA analysis (Fig. 4f) (Table S2). Among these proteins, we found the main collagen proteins (COL1A1 and COL3A1) and other extracellular matrix proteins including fibrillin 1 and 2 (FBN1-2) (Fig. 4g), whose expression profile closely resembled the ones of key enzymes (Fig. 4c) and downstream effectors (Fig. 4d, e, S3c) of the polyamines pathway. To functionally demonstrate the relevance of the polyamines pathway for small intestine regeneration *in vivo*, we used Odc1^loxp/loxp^:Villin-CreERT2 mice. In this model, *Odc1*, the rate-limiting enzyme for polyamine synthesis, is ablated in epithelial cells following tamoxifen administration (Fig. S3e). Thus, we injected 5-FU in young Odc1^fl/fl^:Villin-CreERT2 mice and littermate controls and followed their body weight for 7 days. We found that knockout of *Odc1* alters the body weight profiles in 5-FU injected mice (Fig. 4h), but not in control, PBS-injected mice (Fig. S3f).

**Figure 4.**
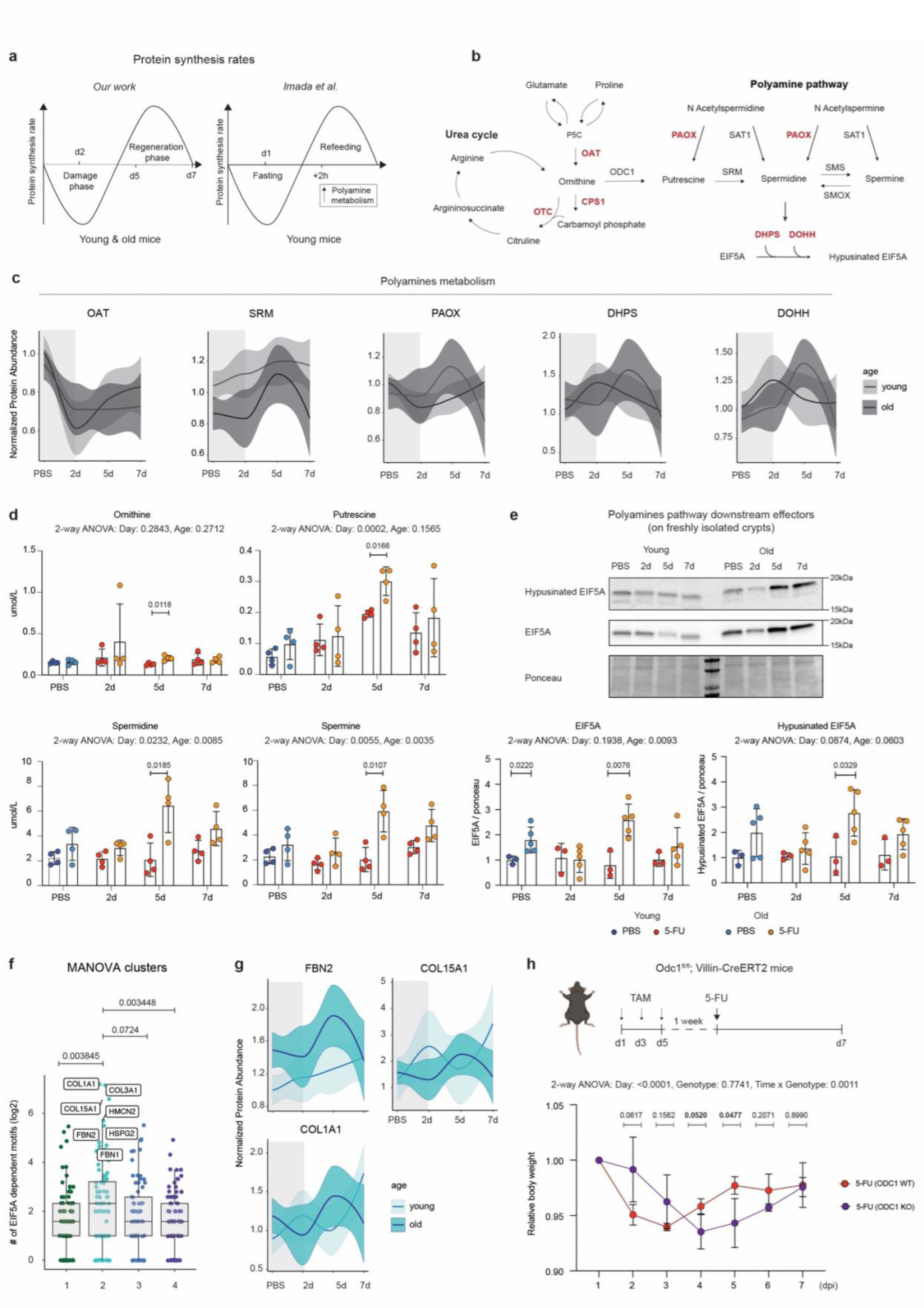
Elevated levels of polyamines upon 5-FU in the aged intestinal epithelium. **a** Schematic representation of dynamics of protein synthesis comparing our damage/regeneration model to the fasting/re-feeding model from Imada *et al.* 2024. **b** Schematic representation of the polyamine biosynthesis and ornithine metabolism pathways. Red proteins are indicating the rate limiting enzymes in these pathways. **c** Protein abundance profiles of the enzymes involved in polyamines biosynthetic pathways. **d** LC-MS quantification of polyamines from crypts lysate. Each dot represents one mouse. *n*=4 mice per condition. Error bars represent the SD. *P*-value was calculated using Welch’s t-test for time point comparison and two-way ANOVA for overall day and age comparisons. **e** Upper: representative immunoblot for hypusinated EIF5A and total EIF5A proteins. Lower: quantification of hypusinated EIF5A and total EIF5A normalized to ponceau staining. Each dot represents one mouse. *n*=3-5 mice per each indicated condition. Error bars represent the SD. *P*-value was calculated using Welch’s t-test for time point comparison and two-way ANOVA for overall day and age comparisons. **f** Distribution of proteins enriched in hypusinated EF5A-dependent motifs across clusters that show distinct dynamics in young and old mice (see Fig. 2e for cluster profiles). Highlighted are proteins that contain the highest number of hypusinated EIF5A-dependent motifs. *P*-value was calculated using Wilcoxon rank sum test. **g** Protein abundance profiles of the COL1A1, COL15A1 and FBN2 proteins in tissue lysate from indicated groups. **h** Upper: Schematic of ODC1 KO induction and 5-FU treatment. Lower: Relative body weight of young ODC1 KO and WT mice treated with a single dose of 5-FU. Body weight of each mouse was normalized to its body weight at the day of injection. *n*=4 ODC1 KO, *n*=3 ODC1 WT mice. Error bars represent the SD. *P*-value was calculated using Welch’s t-test for time point comparison and two-way ANOVA for overall day, treatment and day x treatment comparisons.

**Figure S3.**
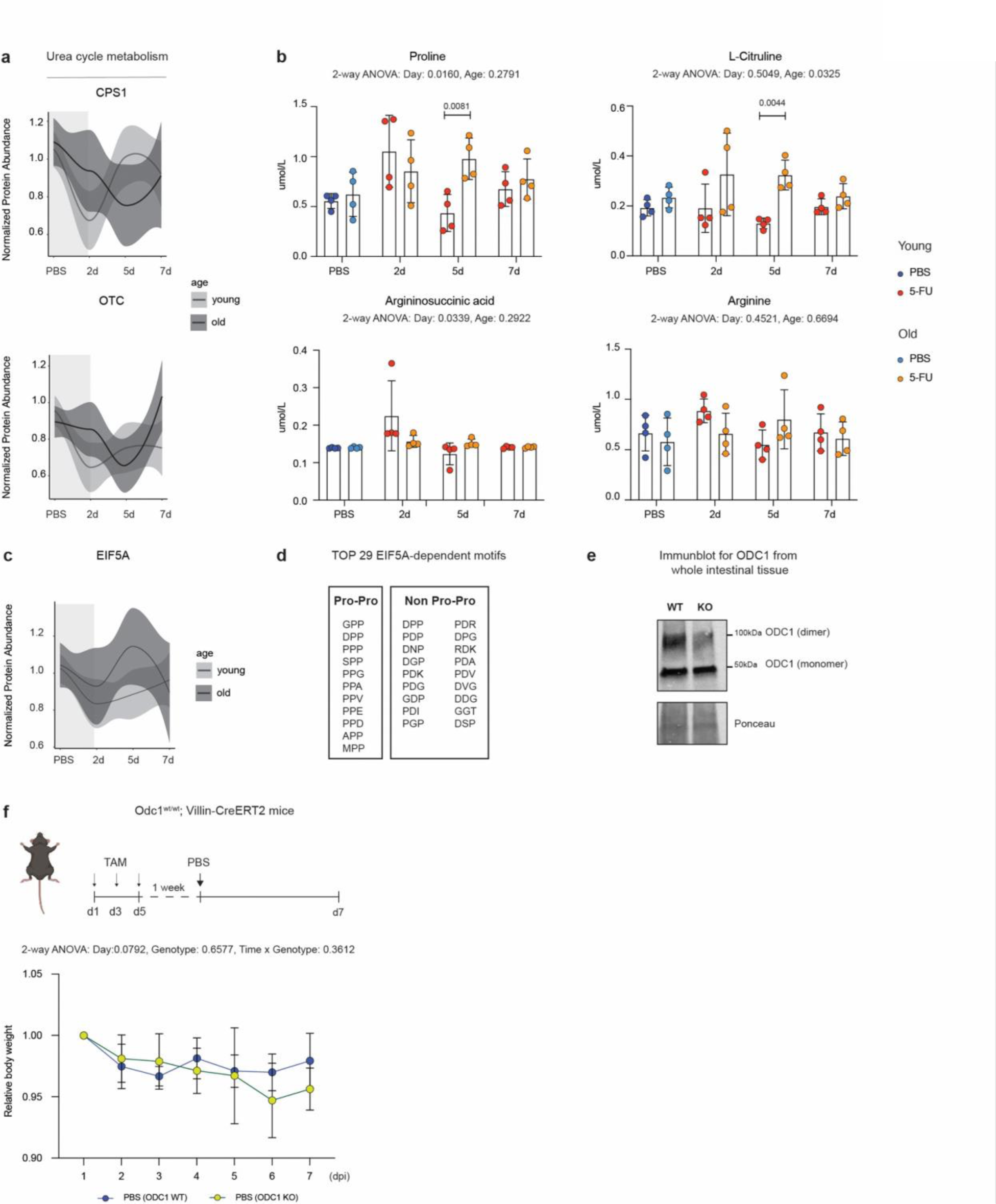
Levels of polyamines during regeneration in young and old intestinal crypts. **a** Protein abundance profiles of the enzymes involved in urea cycle metabolism. **b** Quantification of amino acids by LC-MS. Each dot represents one mouse. *n*=4 mice per condition. Error bars represent the SD. *P*-value was calculated using Welch’s t-test for time point comparison and two-way ANOVA for overall day and age comparisons. **c** Protein abundance profiles of the Eukaryotic Translation Initiation Factor 5A (EIF5A) **d** Top 29 motifs that depend on hypusinated EIF5A for efficient translation according to ^20^. **e** Immunoblot for ODC1. Proteins were extracted from formalin-fixed paraffin embedded (FFPE) whole intestinal tissue samples. Bands corresponding to the dimeric (catalytically active form ^45^) and monomeric forms of ODC1 are highlighted. **f** Upper: Schematic of ODC1 KO induction and PBS treatment. Lower: Relative body weight of young ODC1 KO and WT mice treated with a single dose of 5-FU. Body weight of each mouse was normalized to its body weight at the day of injection. *n*=3 ODC1 KO, *n*=2 ODC1 WT mice. Black line indicates the mean with SD for each day in each treatment group. *P*-value is calculated by two-way ANOVA for overall day, treatment and day x treatment comparisons.

Together, these data demonstrate that the polyamine pathway is induced in response to 5-FU in old epithelia likely to sustain proteostasis and the synthesis of key proteins required for intestinal regeneration, such as collagens and other extracellular matrix components. Additionally, knockout of *Odc1* is sufficient to influence the body-weight recovery following 5-FU also in young mice, further highlighting the role of this pathway for intestinal epithelium regeneration.

### Impact of re-feeding on the regenerative capacity of old intestines

The increased levels of polyamines at 5 dpi, concomitant with the emergence of proteostasis stress, made us hypothesize that the induction of polyamines might be part of an adaptive response of old epithelial cells to injury. We reasoned that interventions that elevate polyamine levels might restore the regenerative capacity of the old intestine to a youthful state. In support of this, it has been recently shown that increased levels of polyamines following post-fasting re-feeding enhance the regenerative capacities of intestinal stem cells ^31^, and we have previously shown that dietary restriction (DR) followed by re-feeding (RF) can reverse aging proteome signatures in intestinal crypts ^30^ affecting multiple proteostasis components including EIF5A (Fig. 5a and S4a). Therefore, we tested whether such a dietary intervention (DR+RF) would be sufficient to restore regenerative capacity upon 5-FU in old mice. First, we confirmed that RF following one month of 30% DR increased the levels of both total and hypusinated EIF5A in intestinal crypts (Fig. S4b). Total and hypusinated EIF5A peaked two days after the beginning of re-feeding and declined thereafter, reaching the levels of ad-libitum (AL) fed mice after 9 days (Fig. 5b). Importantly, the levels of hypusinated EIF5A correlated with the concentration of polyamines in intestinal crypts indicating pathway activation induced by re-feeding (Fig. 5c). Based on these data, we subjected both young and old mice to one month of DR followed by two days of RF prior to 5-FU injection (Fig. 5d). Mice fed ad-libitum (AL) or that underwent DR without re-feeding served as control groups. The dietary interventions had the expected effect on the mice’s body weight prior to 5-FU injection (Fig. S4c). When analyzing body weight curves, we noted that both DR and DR+RF prevented body weight loss following 5-FU injection in young mice. However, in old mice, only DR+RF was sufficient to prevent body weight loss (Fig. 5e and S4d). Next, we investigated further the effect of the dietary interventions on polyamines and downstream effectors in old mice following injury. In line with the previous experiment, mice that underwent DR+RF exhibited elevated levels of total and hypusinated EIF5A (Fig. 5f) at 2 dpi. Consistently, *ex-vivo* puromycin incorporation in intestinal crypts showed increased protein synthesis already at 2 dpi in the DR+RF group, while this occurred only at 5 dpi in AL fed and DR mice (Fig. 5g). Finally, immunoblot for pH3 revealed anticipated (already at 2 dpi) and enhanced (at 5 dpi) cell proliferation in the DR+RF group compared to both AL and DR (Fig. 5h). Together, these data highlight that a dietary intervention based on DR followed by re-feeding activates the polyamine pathway, promotes proteostasis and rescues body weight loss, thus accelerating the recovery of old mice following 5-FU injury.

**Figure 5.**
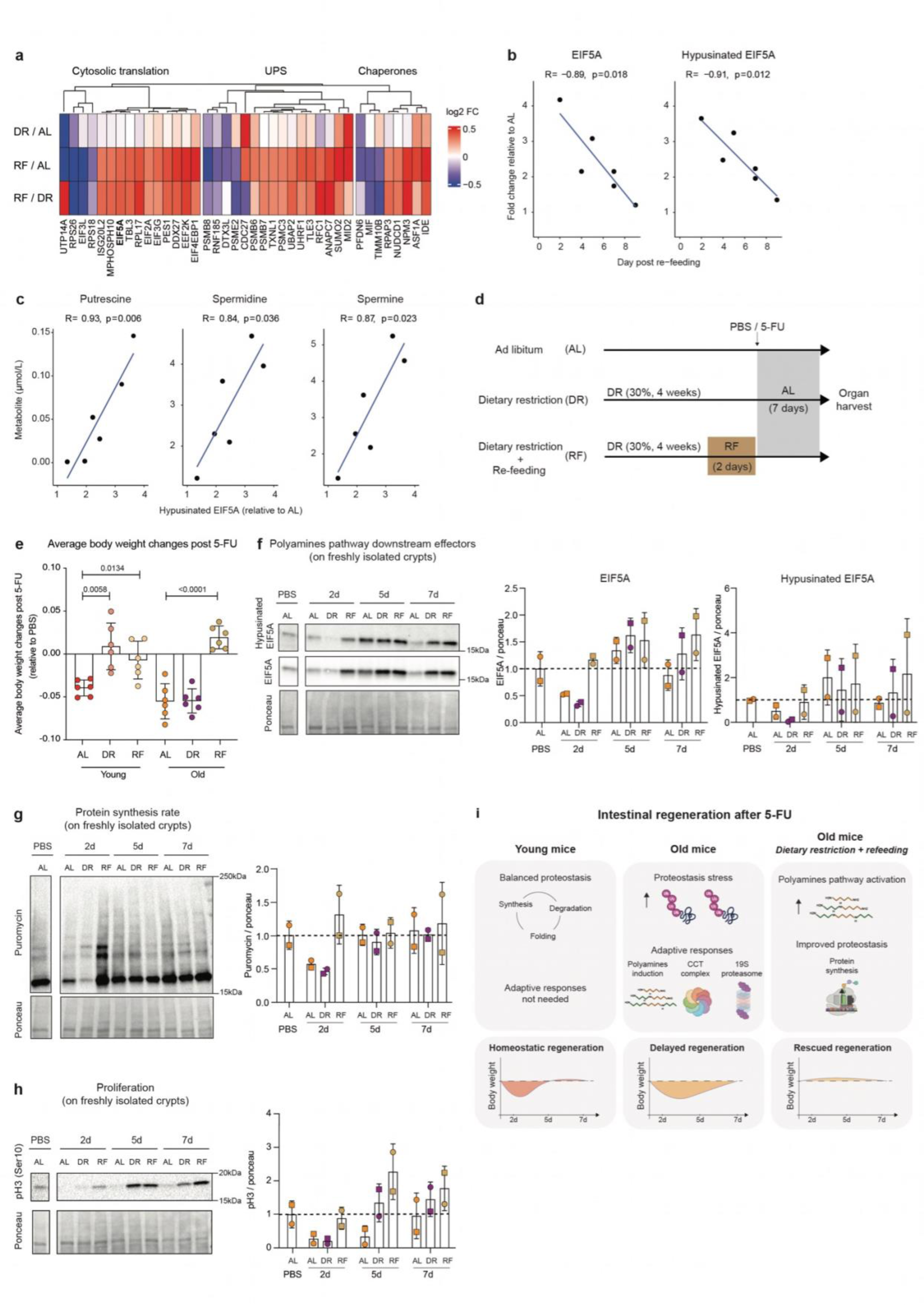
Effect of dietary restriction followed by re-feeding on intestinal epithelium regeneration. **a** Heatmap of fold changes (log2) for significantly affected proteins (absolute log2 fold change > 0.3 and q<0.05) related to proteostasis from aged mice after DR or DR+RF treatments. UPS: ubiquitin proteasome system. Data from Gebert *et al*. 2020 ^30^. **b** Levels of total and hypusinated EIF5A from aged mice at different days after re-feeding, quantified by immunoblot (Fig. S4b) and, **c,** correlation between hypusinated EIF5A and polyamines levels measured on the same intestinal crypt samples from aged mice that underwent DR followed by different days of re-feeding. R values represent Pearson correlation coefficient, P values are the result of the association test between the variables based on Pearson’s product moment correlation coefficient. **d** Scheme of DR and DR+RF treatments following 5-FU induce regeneration. **e** Relative average body weight changes after 5-FU. The body weight of each mouse was normalized to its body weight at the time of injection. For each time point the average body weight of the 5-FU treated group was subtracted from the average body weight of the PBS one. Each dot represents the daily average body weight loss after 5-FU, n= 6. (Averages obtained from *n*=16 PBS, *n*=15 5-FU AL, *n*=4 PBS, *n*=6 5-FU DR, *n*=5 PBS, *n*=7 5-FU RF young mice and *n*=17 PBS, *n*=15 5-FU AL, *n*=4 PBS, *n*=5 5-FU DR, *n*=5 PBS, *n*=5 5-FU RF old mice). P-value was calculated by Welch’s t-test. **f** Left: representative immunoblot for the hypusinated EIF5A and total EF5A proteins on aged mice in the indicated groups. Right: quantification of the hypusinated EIF5A and total EIF5A proteins in the indicated groups. Each dot represents one mouse. *n*=2 mice per group. **g** Upper: Representative immunoblot for the puromycin incorporated proteins on aged mice in the indicated groups. Lower: quantification of the puromycin incorporated proteins in the indicated groups. Each dot represents one mouse. *n*=2 mice per group. **h** Upper: Representative immunoblot for the pH3 protein on aged mice in the indicated groups. Lower: Quantification of pH3 protein relatively to ponceau staining in the indicated groups. Each dot represents one mouse. *n*=2 mice per group. **i** Model of intestinal regeneration dynamics. Left: in young mice, injury leads to rapid small intestine regeneration and body weight recovery within ∼5 days without proteostasis imbalance. Middle: in old mice, injury induces proteostasis stress, leading to delayed regeneration and body weight recovery (∼day 7). Right: old mice subjected to 30 days of dietary restriction followed by 2 days of re-feeding before injury show enhanced polyamine pathway activation, leading to improved protein synthesis and proteostasis post-injury, enhanced intestinal regeneration, and rescued body weight loss.

**Figure S4.**
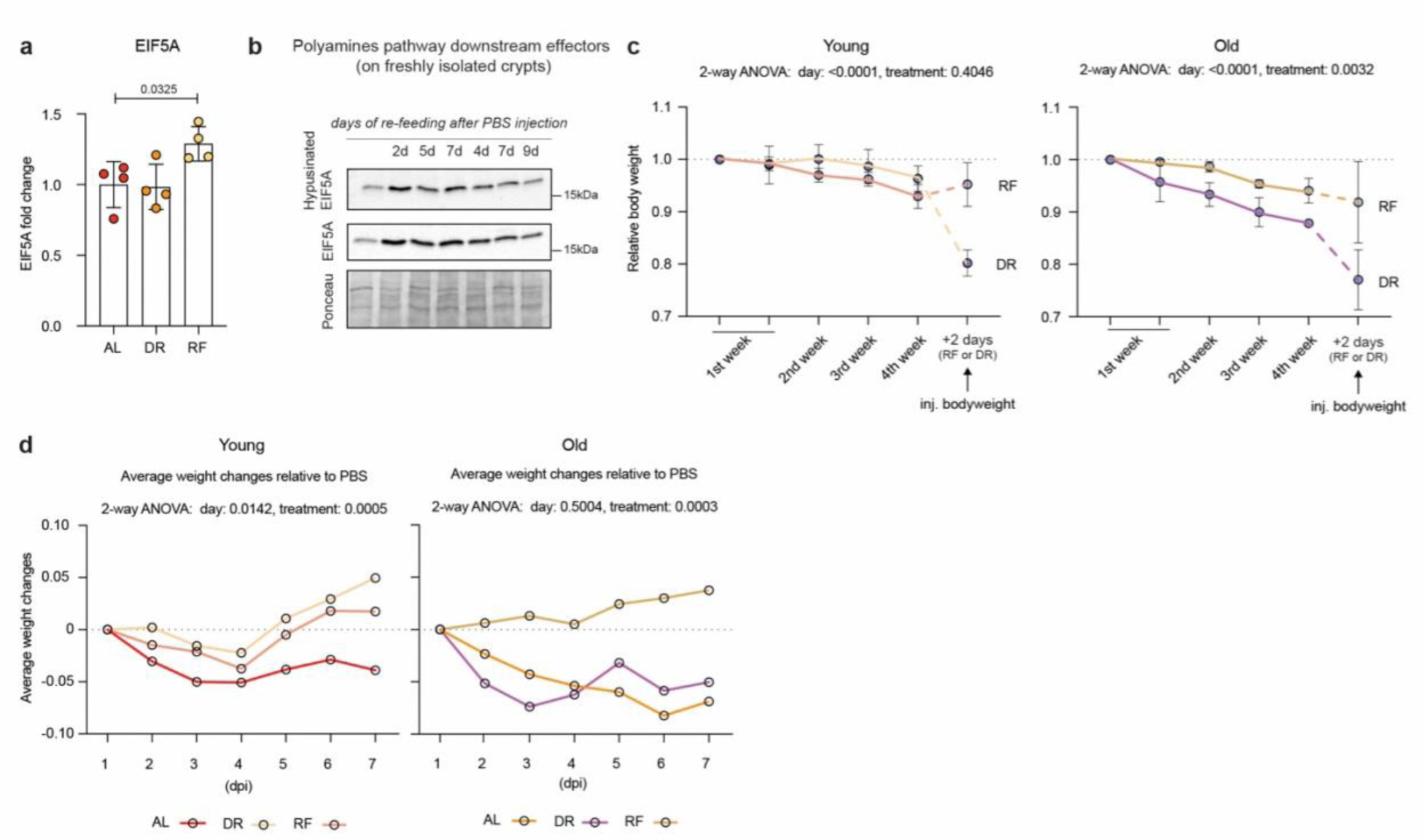
Effect of dietary restriction followed by re-feeding on intestinal epithelium regeneration. **a** Quantification of EIF5A levels in intestinal crypts from old mice that underwent different dietary interventions. Based on proteomics data from Gebert et al. 2020 ^30^. **b** Immunoblot for hypusinated EIF5A and EIF5A proteins on freshly isolated crypts from old mice treated with PBS and re-feed at the indicated time points. **c** Relative body weight of young (left) and old (right) mice under DR for one month followed by two days of re-feeding or continuous DR prior to 5-FU injection. Body weight of each mouse was normalized to its body weight at the time of start. *n*=4 DR, *n*=5 RF young mice and *n*=4 DR, *n*=3 RF old mice. Error bars represent the SD. *P*-value was calculated using two-way ANOVA for overall day and treatment comparisons. **d** Relative body weight of young (left) and old (right) mice from indicated groups after 5-FU injection. Body weight of each mouse was normalized to its body weight at the time of injection. For each time point the average body weight of the 5-FU treated group was subtracted from the average body weight of the PBS one*. (*Averages obtained from *n*=16 PBS, *n*=15 5-FU AL, *n*=4 PBS, *n*=6 5-FU DR, *n*=5 PBS, *n*=7 5-FU RF young mice and *n*=17 PBS, *n*=15 5-FU AL, *n*=4 PBS, *n*=5 5-FU DR, *n*=5 PBS, *n*=5 5-FU RF old mice). *P*-value was calculated using two-way ANOVA for overall day and treatment comparisons.

## Discussion

Multiple studies have indicated that old mice show a delay in the regeneration of intestinal tissue upon different types of injuries compared to young mice ^6,7^. Here, using proteomics, we temporally dissected the phases of intestinal epithelium regeneration and compared protein dynamics in young vs. old mice, uncovering proteomic and metabolomic changes upon damage and regeneration. Intestinal tissue from young mice showed a strong reduction in proliferation 2 days following 5-FU injection, accompanied by a decrease in proteins related to transcription and translation. While the body weight of young mice recovered at day 5 post-injury, old mice required 2 days longer despite showing similar protein synthesis and translation machinery dynamics. The delayed regeneration of old mice was accompanied by the emergence of multiple signs of proteostasis stress exemplified by the accumulation of K48-polyubiquitylated protein at day 5 post-injury.

We established a 5-FU-induced regeneration model in organoid cultures to address the extent to which the delayed regeneration of the old intestine is a cell-intrinsic feature of epithelial cells. We uncovered that intestinal organoids, once challenged with 5-FU, pass through similar regeneration phases as observed *in vivo*, encompassing changes in protein synthesis and proliferative capacity. Additionally, organoids from old mice showed delayed regeneration compared to young ones and, despite some differences, recapitulated proteostasis stress signatures observed *in vivo*, i.e, p62 /SQSTM1 and K48-polyubiquitylated protein accumulation. These data indicate that cell-intrinsic differences in proteostasis might contribute to delayed regeneration of old intestines in addition to changes in niche-derived ^6,7,46^ and systemic factors ^4,5^.

Concomitantly to the emergence of proteostasis stress signatures, we observed that old intestines activate pathways that are typically linked to improved protein homeostasis, such as chaperones and polyamines. We speculate that the activation of these pathways is part of an adaptive response mounted by old intestinal cells to overcome the stress induced by the increase of protein synthesis during post-injury regeneration (Fig. 5i). In line with this hypothesis, we found that proteins containing specific motifs known to depend on hypusinated EIF5A for efficient translation ^20^ cluster together following damage and their abundance profiles precisely follow the dynamics of polyamine pathway activation. Most of these proteins relate to the extracellular matrix, including fibrillin 1 and 2 (FBN1-2) and main collagen proteins such as COL1A1 and COL3A1, whose implication in intestinal tissue repair have been previously documented ^47^. A decline in polyamines has been associated with aging in different tissues, and their supplementation has been shown to be sufficient to increase the median life span of flies, worms, and mice ^19,48,49^. However, our data indicate that polyamines in mice’s small intestines do not decline with age, suggesting that they might not contribute to age-related tissue dysfunction under homeostatic/uninjured conditions.

Emerging evidence shows that polyamines can originate from microbiota or immune cells and then be uptaken by epithelial cells and exert their functions to improve both tissue homeostasis and regeneration ^24,50,51^. Indeed, their supplementation *in vivo* has been shown to promote colonic epithelium regeneration through increased epithelial proliferation and M1/M2 macrophage rebalancing ^24^. Interestingly, recent work found that polyamines are also produced by intestinal stem cells (ISCs) of young mice ^31^. Specifically, polyamines and downstream effectors are induced in ISCs by post-fasting re-feeding, they modulate protein synthesis and, consequently, ameliorate regeneration. However, given their pro-anabolic functions, polyamines were also found to be responsible for increased tumor initiation upon *Apc* deletion ^31^. Together, this evidence suggests the role of the polyamine pathway in supporting protein synthesis in epithelial cells and regeneration following damage. Consistently, we show that knockout of the rate limiting enzymes of polyamine synthesis (ODC1) in epithelial cells is sufficient to alter the response to 5 -FU even in young mice. This indicates that endogenous synthesis of these metabolites by epithelial cells is required also by young epithelial cells, even though we did not detect the same level of induction that we observed in old mice. Further, our data show that treating mice with a dietary intervention that activates the polyamine pathway leads to increased protein synthesis, sustained proliferation, and ultimately prevents the 5-FU induced body weight loss in old mice.

Altogether, our results suggest that activation of the polyamine pathway is one of the underlying mechanisms mediating the beneficial effect of post-DR re-feeding in old mice. Ultimately, given that 5-FU is elective therapy for multiple types of cancers and from the perspective of regenerative medicine, our study provides evidence that modulation of polyamine pathway activation via dietary interventions might represent a valid strategy to promote tissue recovery and alleviate gastrointestinal side effects of 5-FU. However, further studies are necessary to critically evaluate the overall benefits of polyamine pathway activation towards improving regeneration capacity vs. increasing cancer risk in humans.

## Acknowledgments

The authors gratefully acknowledge support from the FLI Core Facilities Proteomics, Imaging, Histology and the Mouse Facility. The authors would like to thank Ellen Späth and Erika Kelmer Sacramento for their support. A.O. is supported by the German Research Council (Deutsche Forschungsgemeinschaft, DFG) via the Research Training Group ProMoAge (GRK 2155), the Else Kröner Fresenius Stiftung (award number: 2019_A79), the Fritz-Thyssen Foundation (award number: 10.20.1.022MN), the Chan Zuckerberg Initiative Neurodegeneration Challenge Network (award numbers: 2020-221617, 2021-230967 and 2022-250618), and the NCL Stiftung. F.N. and A.O. are supported by the German Research Council (Deutsche Forschungsgemeinschaft, DFG) award number NE 2144/5-1. A.T. was supported by an Emmy Noether Award from the German Research Foundation (DFG, 467788900) and the Ministry of Culture and Science of the State of North Rhine-Westphalia (NRW-Nachwuchsgruppenprogramm). A.T. holds the Peter Hans Hofschneider of Molecular Medicine endowed professorship by the Stiftung Experimentelle Biomedizin. F.N. was supported by the AIRC Grant (MFAG 2021 ID 26038) and the Fondazione Ricerca Molinette Grant. The FLI is a member of the Leibniz Association and is financially supported by the Federal Government of Germany and the State of Thuringia.

## Author contributions

Conceptualization: AM, AO, AT, CB, FN, NG, OO, OY

Data curation: AM, AO, AT, CB, JL, MR, OO

Investigation: AM, AT, CB, GA, JR, FC, KS, OO, SY, SK

Methodology: AM, AT, CB, OO, SWM

Project administration: AO, FN

Data analysis: AM, AO, AT, CB, OO

Supervision: AO, AT, FN, RH, OY

Visualization: AM, AO, OO

Writing – original draft: AM, AO, CB, OO

Writing – review & editing: AT, FN

## Methods

### Mice

Young (3–4 months old) and old (21–25 months old) male wildt-ype C57BL6/J mice obtained from Janvier or from internal breeding at the Leibniz Institute on Aging – Fritz Lipmann Institute (FLI). They were group housed and maintained in a Specific Opportunistic Pathogen Free (SOPF) animal facility in Fritz Lipmann Institute with 12 h of light/dark cycle and fed with a standard mouse chow at Temperature 20 ± 2 °C, rlH 55%± 15. Experiments were conducted according to protocols approved by the state government of Thuringia Thüringer Landesamt für Verbraucherschutz (TLV) authority (licenses number: FLI-20-003).

Young (2-4 months) *Odc1^loxp/loxp^; Villin-CreERT2* and *Odc1^wt/wt^; Villin-CreERT2* mice were generated by crossing *Odc1^loxp/loxp^*mice to *Villin-CreERT2* mice under the husbandry care of the Department of Comparative Medicine in Koch Institute for Integrative Cancer Research. Use of 5-FU in Yilmaz lab was approved in the Committee on Animal Care (CAC) protocol at MIT.

### *In vivo* treatments

5-FU treatment: 5-FU (Sigma-Aldrich #F6627) was solubilized in PBS (Sigma-Aldrich #D8537-100ML) at a concentration of 20 mg/ml. Young and old mice were then injected intraperitoneally with 5-FU (150 mg/kg) or PBS as control (single injection in the morning). Mice were sacrificed and intestinal tissue harvested 2,5 and 7 days after injection for downstream analysis. <u>Tamoxifen treatment</u>: Tamoxifen injections were achieved by intraperitoneal (i.p.) injection suspended in sunflower seed oil (Spectrum S1929) at a concentration of 4 or 10 mg/ml. *Odc1^loxp/loxp^; Villin-CreERT2 and Odc1^wt/wt^; Villin-CreERT2 mice* were injected with 2.5 mg/25gr every other day (Day 1, 3 and 5), 5-FU or PBS was then injected 1 week after the last injection.

### Dietary restriction and re-feeding mice experiment

Mice were separated to single cages two weeks prior to the dietary treatment. Food weight and body weight were measured directly after separating the animals and before the dietary treatment. The daily food intake per animal was calculated and used for calculating the amount that refers to 70% for every animal. Food was given in the morning; once per day to the DR and the DR+RF animals for one month of DR period. AL animals had unlimited access to food throughout the experiment. Animals of the DR+RF cohort underwent the same dietary treatment as DR animals and received unlimited access to food for 2 days prior to injection. Mice were injected Intraperitoneally with 5-FU (150 mg/kg) or PBS as control in the morning and unlimited food access provided after injection to all of the cohorts until end of experiments.

### Hematoxylin and Eosin staining

Sections on slides were deparaffinized by xylene twice 5 minutes each and rehydrated by 100%, 90%, 70%, 50% and 30% ethanol for 5 minutes each and washed with milliQ water for 10 minutes. The sections were stained in hematoxylin for 3 minutes, washed in water for 5 minutes and stained in eosin for 3 minutes. The sections were dehydrated through 30%, 50%, 70%, 90% and 100% ethanol and incubated in xylene twice for 5 minutes each. Slides were mounted by using xylene based mounting medium, Depex (VWR #100503-836). Section cutting and staining was done by the core FLI histology facility. Images of stained sections were acquired using Axio Imager (Zeiss, Germany) and analyzed by the ZEN blue software v2 (Zeiss, Germany). For further image analysis, the graphics tools for counting and measuring the ZEN software were used.

### Immunofluorescence on paraffin-embedded tissue sections

5 μm paraffin sections were deparaffinized by two times immersion in xylene (10 minutes each time) and rehydrated by immersion in a series of graded ethanol dilutions 100%, 90%, and 70% for 5 minutes each. Epitope retrieval was performed by preheating the sections 5 minutes at full power microwave (900W) in 10mM sodium citrate buffer pH 6.5 until boiling, followed by 10 minutes at a sub-boiling temperature (600W). The sections were cooled down for 20 minutes, washed in PBS and blocked with5% BSA/PBS for 1 h at RT in a humid chamber. Sections were stained with rabbit anti-phospho-histone H3 (Ser10) antibody (1:100) in 1% BSA/PBS for 16 h at 4 degree in the humid chamber. This was followed by washing in T-PBS (0.1% Tween 20 in PBS v/v, 3 × 5 minutes) and subsequent incubation for 30 minutes with secondary anti-rabbit IgG conjugated with AF594 (1:1000). The slides were then washed in T-PBS 3 × 5 minutes and mounted with a mounting medium including DAPI. Images of stained sections were acquired using Axio Imager (Zeiss, Germany) and analyzed by the ZEN blue software v2 (Zeiss, Germany). For further image analysis, the graphics tools for counting and measuring the ZEN software were used.

### Small intestinal crypts isolation

After euthanizing the mice with CO2, the small intestine was isolated and cleaned with cold PBS. The villi were removed by scraping with glass coverslip and the villi free intestinal pieces (2 cm) were washed with cold PBS and incubated in 5mM EDTA/PBS for 30-40 min at 4 °C on a rotator. The tissue was transferred to a new 50 ml falcon filled with fresh cold PBS and manually shaken for 30 sec. The crypt solution was filtered using a 70 μm cell strainer (Corning #431751) and centrifuged at 450 × g for 5 min at 4 °C. Isolated crypts were immediately used or snap-frozen in liquid nitrogen and stored at −80 °C for further experiments.

### Organoids culture and *in vitro* 5-FU treatment

Small intestine crypts were isolated from mice, as described above. They were then embedded in 70% v/v Matrigel Growth Factor Reduced (Corning #356231) mixed with organoid medium. Organoid medium consists of Advanced DMEM F12 (Life Technologies #12634-010) supplemented with Noggin 100 ng/ mL (Peprotech #250-38), recombinant epidermal growth factor 50 ng/mL (Peprotech #315-09), R-spondin 1 mg/mL (home-made), N2 1X (Life Technologies #17502048) and B27 1X (Life Technologies #12587010), 1X GlutaMax (Thermo Fisher scientific #35050061), 10mM HEPES (Thermo Fisher scientific #15630080), 0.5 U/ml penicillin/streptomycin (Thermo Fisher scientific #15140122). Next, crypts were plated onto a flat bottom 24-well plate (VWR #734-2325) for 10 minutes in a 37°C incubator with 5% CO2 until Matrigel solidifies. Matrigel-crypts domes were then overlaid with 500 mL organoid medium, which was changed every three days. For in vitro 5-FU treatment, organoids were let grow for a week after initial passage and then treated with the reported concentrations of 5-FU solubilized in PBS at a concentration of 20 mg/ml. 24 hours after, the organoids were passed 1:1 into new plates and collected/analyzed at the indicated time points.

### Protein lysate and Immunoblotting

For protein lysate, intestinal crypts or organoids were lysed in RIPA buffer (RIPA buffer (150 mM Sodium Chloride (Roth #P029.2), 1 % Triton X-100 (v/v Roth #3051.3), 0.5 % Sodium Deoxycholate (Thermo Fisher #89904), 0.1 % SDS (w/v Sigma Aldrich #75746-250G), 50 mM Tris (Roth #4855.2) pH8) supplemented with cOmplete™, Mini, EDTA-free Protease Inhibitor (Roche #11836170001) and with PhosSTOP™ Phosphatase Inhibitor (Roche #4906837001). Samples were then sonicated in a Bioruptor Plus (Diagenode, Belgium) for 10 cycles with 1 min ON and 30 s OFF with high intensity at 4 °C and then centrifuged max speed for 5 min. The resulting supernatant was then collected, and the proteins quantified using Pierce™ BCA protein assay kit (Thermo Scientific #23225) following manufacturer’s instructions. Protein lysates were then stored at -80°C before use. For immunoblotting 15 µg of protein lysate in 4× loading buffer (1.5 M Tris pH 6.8, 20% (w/v) SDS, 85% (v/v) glycerin, 5% (v/v) β-mercaptoethanol) were denatured at 95°C and then separated via SDS-PAGE on the 4-20% Mini-Protean TGX Gels (Bio-Rad #4561096). For molecular weight estimation, 5 µl of Precision Plus Protein™ Dual Color standard (Bio-Rad #1610374) was applied. Proteins were then transferred using a semi-wet transfer approach into Nitrocellulose blotting membranes (Carl Roth #200H.1) using the Trans-Blot® Turbo™ Transfer Starter System (Bio-Rad Laboratories, CA, USA). Membranes were then stained with Ponceau S (Sigma #P7170-1L) for 5 min on a shaker (Heidolph Duomax 1030), washed with Milli-Q water, imaged on a Molecular Imager ChemiDocTM XRS + Imaging system (Bio-Rad Laboratories CA, USA) and destained by 2 washes with PBS and 2 washes in TBST (Tris-buffered saline (TBS, 25 mM Tris, 75 mM NaCl), with 0.5% (v/v) Tween-20) for 5 min. After incubation for 5 min in EveryBlot blocking buffer (Bio-Rad Laboratories #12010020), membranes were incubated overnight with primary antibodies: Mouse-Puromycin (Millipore Sigma #MABE343), Rabbit-Ubiquitin linkage specific K48 (Abcam #ab140601), Rabbit-SQSTM1/p62 (Abcam #ab91526), Mouse-S6 ribosomal protein (Cell Signaling Technologies #2317), Mouse-Ubiquitin P4D1 (Santa Cruz Biotechnology #sc-8017), Rabbit-Cleaved Caspase 3 (Cell Signaling Technologies #9661), Rabbit-Caspase 3 (Cell Signaling Technologies #9662), Rabbit-phospho H3 (Cell Signaling Technologies # 9701), Rabbit-Hypusinated EIF5A (Millipore Sigma #ABS1064), Mouse-EIF5a (BD Biosciences #611977), Rabbit-ODC1 (Abcam #ab137679) diluted (1:1000) in enzyme dilution buffer (0.2% (w/v) BSA, 0.1% (v/v) Tween20 in PBS) at 4 °C on a tube roller (BioCote® Stuart® SRT6). Membranes were then washed 3 times with TBST for 10 min at room temperature and incubated with horseradish peroxidase coupled secondary antibodies (Dako anti-rabbit #P0448 or anti-mouse #P0447) at room temperature for 1 h (1:2000) in enzyme dilution buffer. After 3 more washes for 10 min in TBST, chemiluminescent signals were detected using ECL (enhanced chemiluminescence) Pierce ECL detection kit (Thermo Fisher Scientific #32109). Signals were acquired on the Molecular Imager ChemiDocTM XRS + Imaging system and analyzed using the Image Lab 6.1 software (Bio-Rad laboratories, CA, USA). When needed, membranes were stripped using Restore Western Blot Stripping Buffer (Thermo Scientific #21059) 10ml x 30 min, washed 3 times with TBST, blocked, and then incubated with the second primary antibody.

### Immunoblot from formalin-fixed paraffin embedded (FFPE) whole intestinal tissue samples

Sections on slides were deparaffinized by xylene twice 5 minutes each and rehydrated by 100%, 90%, 70%, 50% and 30% ethanol for 5 minutes each and washed with milliQ water for 10 minutes. After deparaffinization, the sections were gently scraped with a scalpel and transferred to a PCR tube with 80ul of protein lysis buffer (DTT 1M, Tris 1M pH 8.0, SDS 20% in milliQ water) and 4× loading buffer (1.5 M Tris pH 6.8, 20% (w/v) SDS, 85% (v/v) glycerin, 5% (v/v) β-mercaptoethanol). Samples were then sonicated in a Bioruptor Plus (Diagenode, Belgium) for 15 cycles with 1 min ON and 30 s OFF at room temperature and then boiled for 1 hour to de-crosslink proteins. The sonication and boiling steps were repeated twice. Resulting protein lysate was then used for Immunoblot as previously reported.

### Protein synthesis assay

To assess protein synthesis rate, we used puromycin which incorporates at the C-terminus of nascent polypeptide chains. Freshly isolated crypts were incubated at 37°C with organoids medium including 10 ug ml-1 puromycin for 15 min, centrifuged for 5 min, and then, resulting pellet was eluted with RIPA buffer containing cOmplete™, Mini, EDTA-free Protease Inhibitor (Roche #11836170001) and with PhosSTOP™ Phosphatase Inhibitor (Roche #4906837001). For *in vitro* protein synthesis assay, intestinal organoids were collected in PBS and centrifuged for 5 min, the resulting pellet was then incubated with organoids medium supplemented with 10 ug ml-1 puromycin (Millipore Sigma #P8833) for 15 min, centrifuged 5 min and eluted with RIPA buffer supplemented with proteases and phosphatases inhibitors. The up-taken puromycin to the nascent polypeptide was then analyzed with immunoblotting using mouse monoclonal anti-puromycin antibody (Millipore Sigma #MABE343) diluted 1:1000.

### Quantitative real-time PCR

After sacrificing the mice using CO2, intestinal tissue was immediately placed in QIAzol Lysis Reagent (Qiagen #79306) and stored at −80 °C for RNA isolation accordingly. 1 μg of total RNA was used to prepare cDNA using iScript cDNA Synthesis Kit (Bio-Rad Laboratories #1708891) according to the manufacturer’s protocol. Quantitative real-time PCR analysis was performed on a CFX384 Touch Real-Time PCR System (Bio-Rad Laboratories, CA, USA) using SYBR GreenER qPCR SuperMix (Thermo Fisher Scientific #11761500). Each reaction was performed in a 9 μl qPCRmix and 1 μl of 1∶10 diluted cDNA. qRT-PCR conditions were 3 min at 95 °C, then 45 cycles of 15 s at 95 °C and 30 s at 61 °C. To obtain amplicon data, a melting curve analysis was performed after each PCR run. Each sample was analyzed in triplicate. The gene expression was determined following the delta Ct method and normalized to Beta-actin. The following mouse primers were used: *Actb* forward 5′-TCTTTGCAGCTCCTTCGTTG -3′, reverse 5′-ACGATGGAGGGGAATACAGC-3′; *p62* forward 5′-AGGATGGGGACTTGGTTGC-3′, reverse 5′-TCACAGATCACATTGGGGTGC-3′.

### Sample preparation for mass spectrometry (MS) analysis

3cm of intestinal tissues from distal ileum were thawed and transferred into Precellys® lysing kit tubes (Keramik-kit 1.4/2.8 mm, 2 ml (CKM)) containing 500 μl of PBS supplemented with 1 tab of cOmplete™, Mini, EDTA-free Protease Inhibitor per 50 ml. For homogenization, tissues were shaken twice at 6000 rpm for 30 s using Precellys® 24 Dual (Bertin Instruments, France) and the homogenate was transferred to new 1.5 ml Eppendorf tubes. Based on estimated protein content, 100 µg of protein was processed for further analyses. Volumes were adjusted using PBS and one volume equivalent of 2× lysis buffer (100 mM HEPES pH 8.0, 2% (w/v) SDS) was added. Samples were sonicated in a Bioruptor Plus (Diagenode, Belgium) for 10 cycles with 1 min ON and 30 s OFF with high intensity at 20 °C. Samples were heated for 10 min at 95°C and a second sonication cycle was performed as described above. Samples were reduced using 10 mM Dithiothreitol (DTT) (Roth #6908.3) for 30 min at room temperature and alkylated using freshly made 15 mM iodoacetamide (IAA) (Sigma-Aldrich #I1149) for 30 min at room temperature in the dark. Subsequently, proteins were acetone precipitated and digested using LysC (Wako sequencing grade #125-05061) and trypsin (Promega sequencing grade #V5111), as described by Buczak et al. ^52^. The digested proteins were then acidified with 10% (v/v) trifluoracetic acid and desalted using Waters Oasis® HLB µElution Plate 30 µm (Waters #186001828BA) following manufacturer instructions. The eluates were dried down using a vacuum concentrator and reconstituted in 5% (v/v) acetonitrile, 0.1% (v/v) formic acid. Samples were transferred to an MS vial, diluted to a concentration of 1 µg/µl, and spiked with iRT kit peptides (Biognosys AG #Ki-3002-2) prior to analysis by LC-MS/MS.

### Proteomics data acquisition

Peptides were separated in trap/elute mode using the nanoAcquity MClass Ultra-High Performance Liquid Chromatography system (Waters, MA, USA) equipped with trapping (nanoAcquity Symmetry C18, 5 μm, 180 μm × 20 mm) and an analytical column (nanoAcquity BEH C18, 1.7 μm, 75 μm × 250 mm). Solvent A was water and 0.1% formic acid, and solvent B was acetonitrile and 0.1% formic acid. 1 μl of the samples (∼1 μg on column) were loaded with a constant flow of solvent A at 5 μl/min onto the trapping column. Trapping time was 6 min. Peptides were eluted via the analytical column with a constant flow of 0.3 μl/min. During the elution, the percentage of solvent B increased nonlinearly from 0–40% in 120 min. The total run time was 145 min, including equilibration and conditioning. The LC was coupled to an Orbitrap Exploris 480 (Thermo Fisher Scientific, Germany) using the Proxeon nanospray source. The peptides were introduced into the mass spectrometer via a Pico-Tip Emitter 360-μm outer diameter × 20-μm inner diameter, 10-μm tip (New Objective) heated at 300 °C, and a spray voltage of 2.2 kV was applied. The capillary temperature was set at 300°C. The radio frequency ion funnel was set to 30%. For DIA data acquisition, full scan mass spectrometry (MS) spectra with a mass range 350– 1650 m/z were acquired in profile mode in the Orbitrap with the resolution of 120,000 FWHM. The default charge state was set to 3+. The filling time was set at a maximum of 60 ms with a limitation of 3 × 10^6^ ions. DIA scans were acquired with 40 mass window segments of differing widths across the MS1 mass range. Higher collisional dissociation fragmentation (stepped normalized collision energy; 25, 27.5, and 30%) was applied, and MS/MS spectra were acquired with a resolution of 30,000 FWHM with a fixed first mass of 200 m/z after accumulation of 3 × 106 ions or after filling time of 35 ms (whichever occurred first). Data were acquired in profile mode. For data acquisition and processing of the raw data, Xcalibur 4.3 (Thermo Fisher Scientific, Germany) and Tune version 2.0 were used.

### Proteomics data analysis

DIA were searched against a spectral library generated using Spectronaut Professional v15.4 (Biognosys AG, Switzerland). For library creation, the DDA and DIA raw files were searched with Pulsar (Biognosys AG, Switzerland) against the mouse UniProt database (*Mus musculus,* v. 160106, 16,747 entries) with a list of common contaminants (247 entries) appended, using default settings. The library contained 101920 precursors and 6569 protein groups satisfying an FDR of 1% at both the precursor and protein group level.

DIA data were searched against this spectral library using Spectronaut Professional (v.16.0) and default settings. Protein quantification was performed in Spectronaut using default settings, except: Protein LFQ method= QUANT 2.0; Proteotypicity Filter = Only protein group specific; Major Group Quantity = median peptide quantity; Major Group Top N = OFF; Minor Group Quantity = median precursor quantity; Minor Group Top N = OFF; Data Filtering = Q value sparse; Normalization Strategy = Local normalization; Row Selection = Automatic. The data (report table) was then exported, and further data analyses and visualization were performed with Rstudio using in-house pipelines and scripts.

### Clustering and comparison of young and old protein abundance profiles

Normalized protein quantities for the regeneration time course were filtered for protein groups quantified by at least two unique (proteotypic) peptides. All the time points post 5-FU (2d, 5d and 7d) were compared to the median of PBS injected control samples, and protein groups that showed an absolute log2 fold change > 0.5 at any time point were selected for clustering analysis. Determination of an optimal number of clusters, clustering, and REACTOME pathway enrichment analysis for each cluster was performed with ClueR ^53^ using the log2 fold change to PBS as input. For the comparison of young and old mice, ratios to PBS control from individual replicates were log2 transformed and missing values removed. Protein abundance profiles were compared using Multivariate ANalysis Of VAriance (MANOVA) by setting protein ratios at different time points as dependent variables and age group as treatment.

### Polyamines measurement

#### Chemicals

Ultrapure and desalted water with a resistivity of 18.2 M Ω/cm was generated by a Sartorius Stedim water purification system (Sartorius, Germany). All the chemicals used for polyamines measurement are shown in Key resource table - Chemicals used for polyamines measurement.

#### Extraction of Amino Acids and Polyamines

Polyamines were extracted by a two-step liquid extraction adapted from Sellik et al. 2011^54^. Briefly, 50k crypts were snap-frozen in liquid nitrogen for quenching. The frozen crypts were extracted by adding 500 µL prechilled methanol (-80 °C), 20 µL of internal standard (L-arginine- ^13^C_6_, spermidine-^2^H_6_, spermine -^2^H_8_ at a concentration of 100 µM which will result in a concentration of 10 µM in the final extract) and vortexed for 2 minutes until no cell clumps were visible. Afterwards, the samples were sonicated in a chilled (0 °C) ultrasonic bath for two minutes. The cells were kept at -80 °C for 5 minutes and subsequently thawed. The thawed cells were vortexed and sonicated for 2 minutes and centrifuged at 3000 g for 5 minutes. The supernatant was collected and 250 µL of water acidified with 0.1 % acetic acid (LC-MS grade) was added. The samples were vortexed and sonicated for 2 minutes and subject to the same freezing-thawing cycle described before. The thaw samples were again vortex and sonicated for 2 minutes, respectively. Next, the samples were centrifuged at 3000g for 5 minutes and the supernatant was collected. The combined supernatant was dried in a vacuum-centrifuge for 45 min until complete dryness. The dried extract is resuspended in 200 µL acetonitrile/water (50/50; v/v) by sonicating and vortexing for 2 minutes, respectively. The final extract was obtained as the supernatant following a final centrifugation at 12.000 rpm for 2 minutes.

#### Dansylation of Polyamines

Dansylation was carried out by using 80 µL of the obtained extract or of a standard solution with a known concentration. The solution is dried and then reconstituted in 80 µL water, 40 µL PBS (1x), and 20 µL of 1 M NaOH. The reaction was started by adding 80 µL of a dansylchloride solution (10 mg/mL in acetone). The mixture was incubated at 55 °C for 30 minutes. Afterward, the solution was dried in a vacuum centrifuge to complete dryness for 45 minutes. The residue was dissolved in 80 µL of acetonitrile/water (50/50; v/v) and subsequently analyzed by LC-MS.

#### LC-MS analysis and quantification

For polyamine analysis, an Agilent 1290 Infinity LC system coupled to an Agilent 6470 QqQ-MS was used (Agilent Technologies, Germany). Liquid chromatographic (LC) separation was carried out using a Zorbax Eclipse Plus C18 RRHD (50 x 2.1 mm, 1.8 µm; Agilent Technologies Inc., Waldbronn, Germany). The elution was carried out by utilizing a gradient at a flow rate of 400 μL/min with water (10 mM ammonium formate pH 3.5) as solvent A and acetonitrile/water (90/10; v/v; 10 mM ammonium formate pH 3.5) as solvent B. The linear gradient was: 0 min, 15% B; 2 min, 15% B; 7 min, 100% B; 10 min, 100% B followed by 2 min at initial condition for re-equilibration. Column temperature was 45 °C, injection volume 10 µL. Ionization was carried out in ESI positive mode by using the Agilent jet stream source. The following MS parameters were used: capillary 4500 V, nozzle voltage 1000 V, gas temp. 275 °C, gas flow 10 L/min, nebulizer gas pressure 25 psi, sheath gas temp. 350 °C, sheath gas flow 10 L/min. Detection was carried out in selected reaction monitoring (SRM) using the following optimized transitions for the transitions and parameters shown in Table 1. For amino acid analysis, the same system was used. LC separation was carried out using a AdvanceBio MS Spent Media HILIC column (2.1x150 mm 2.7 µm; Agilent Technologies, Germany). For elution a binary gradient with water 10 mM ammonium formate + 0.1% formic acid as A and 90/10 ACN/Water 10 mM ammonium formate + 0.1% formic acid as B was used. The flow rate was 400 µL and the linear gradient program was: 0 min 90% B, 2 min 90% B, 12 min 40% B, and 12.1 min 5% B, which was held for 6 min. followed by 4 min at initial conditions for re-equilibration. Column temperature was 45 °C, injection volume 10 µL. For ESI and MS detection the following parameters were used: capillary 3000 V, nozzle voltage 0 V, gas temp. 225 °C, gas flow 6 L/min, nebulizer gas pressure 40 psi, sheath gas temp. 225 °C, sheath gas flow 10 L/min. Detection was carried out in selected reaction monitoring (SRM) using the following optimized transitions for the transitions and parameters shown in Table 2. Non-derivatized polyamines were only monitored with this method but spermidine and spermine showed rather high limit-of-detections. Therefore, quantification of these compounds was done in a separate method.

**Table 1:** Transition settings and internal standards for the analysis of polyamines.

| Compound | Internal standard | *m/z* Q1 | *m/z* Q2 | Dwell time (ms) | Fragmentor (V) | Collision Energy (V) | Cell Accelerator (V) |
| --- | --- | --- | --- | --- | --- | --- | --- |
| Dansylchloride | - | 234.7 | 234.7 | 50 | 200 | 0 | 4 |
| L-Arginine-<sup>13</sup>C<sub>6</sub> | - | 408.1 | 70.1 | 50 | 200 | 35 | 4 |
| N1-Ac-Putrescine | Spermidine-<sup>2</sup>H<sub>6</sub> | 364.1 | 170.1 | 50 | 160 | 30 | 4 |
| N1-Ac-Spermidine | Spermidine-<sup>2</sup>H<sub>6</sub> | 654.1 | 101.1 | 50 | 200 | 40 | 2 |
| N1-Ac-Spermine | Spermidine-<sup>2</sup>H<sub>6</sub> | 944.1 | 360.1 | 50 | 200 | 53 | 2 |
| Putrescine | Spermidine-<sup>2</sup>H<sub>6</sub> | 555.2 | 170.1 | 50 | 220 | 40 | 3 |
| Spermidine | Spermidine-<sup>2</sup>H<sub>6</sub> | 845.4 | 360.1 | 50 | 220 | 46 | 2 |
| Spermidine-<sup>2</sup>H<sub>6</sub> | - | 851.4 | 366.1 | 50 | 220 | 43 | 2 |
| Spermine | Spermine-<sup>2</sup>H<sub>8</sub> | 1135.4 | 360.1 | 50 | 220 | 50 | 3 |
| Spermine-<sup>2</sup>H<sub>8</sub> | - | 1143.4 | 367.1 | 50 | 180 | 55 | 3 |

**Table 2:** Transition settings and internal standards for the analysis of amino acids.

| Compound | Internal standard | m/z Q1 | m/z Q2 | Dwell time (ms) | Fragmentor (V) | Collision Energy (V) | Cell Accelerator (V) |
| --- | --- | --- | --- | --- | --- | --- | --- |
| Argininosuccinic acid | L-Arginine-<sup>13</sup>C<sub>6</sub> | 291.1 | 70.2 | 50 | 195 | 25 | 6 |
| Cadaverin | Spermidine-<sup>2</sup>H<sub>6</sub> | 103.1 | 86.1 | 50 | 135 | 13 | 4 |
| L-Anserine | L-Arginine-<sup>13</sup>C<sub>6</sub> | 241.1 | 109.0 | 50 | 135 | 25 | 4 |
| L-Arginine | L-Arginine-<sup>13</sup>C<sub>6</sub> | 175.1 | 70.2 | 50 | 150 | 25 | 6 |
| L-Arginine-<sup>13</sup>C<sub>6</sub> | - | 181.1 | 74.2 | 50 | 135 | 25 | 4 |
| L-Aspartic acid | L-Arginine-<sup>13</sup>C<sub>6</sub> | 134.0 | 88.0 | 50 | 120 | 17 | 3 |
| L-Citrulline | L-Arginine-<sup>13</sup>C<sub>6</sub> | 176.1 | 70.2 | 50 | 145 | 25 | 6 |
| L-Ornithine | L-Arginine-<sup>13</sup>C<sub>6</sub> | 133.1 | 70.2 | 50 | 130 | 29 | 4 |
| N1,N12-Diacetylspermine | Spermidine-<sup>2</sup>H<sub>6</sub> | 287.2 | 100.1 | 50 | 135 | 20 | 4 |
| N1-Ac-Putrescine | Spermidine-<sup>2</sup>H<sub>6</sub> | 131.1 | 72.1 | 50 | 135 | 15 | 4 |
| N1-Ac-Spermidine | Spermidine-<sup>2</sup>H<sub>6</sub> | 188.2 | 72.1 | 50 | 160 | 20 | 4 |
| N1-Ac-Spermine | Spermidine-<sup>2</sup>H<sub>6</sub> | 245.2 | 112.2 | 50 | 145 | 25 | 4 |
| Proline | L-Arginine-<sup>13</sup>C<sub>6</sub> | 116.1 | 70.1 | 50 | 135 | 15 | 4 |
| Putrescine | Spermidine-<sup>2</sup>H<sub>6</sub> | 89.1 | 72.2 | 50 | 110 | 11 | 4 |
| Spermidine | Spermidine-<sup>2</sup>H<sub>6</sub> | 146.2 | 72.1 | 50 | 135 | 16 | 5 |
| Spermidine-<sup>2</sup>H<sub>6</sub> |  |  |  |  |  |  |  |
| Spermine | Spermine-<sup>2</sup>H<sub>8</sub> | 203.2 | 112.1 | 50 | 145 | 18 | 6 |
| Spermine-<sup>2</sup>H<sub>8</sub> | - | 211.3 | 120.1 | 50 | 145 | 18 | 6 |

Quantification in crypt cells was performed by an external calibration in a range from 0.1 µM to 90 µM using 3 deuterated internal standards (L-arginine-^13^C_6_, spermidine-^2^H_6,_ and spermine-^2^H_8_) at a concentration of 10 µM. For calibration, the analyte to internal standard area ratios were linearly fitted to the corresponding concentration ratios and compared to the area ratios detected in the samples.

### Key Resource Table

#### Experimental Models: Organisms/Strains

|  |  |  |
| --- | --- | --- |
| *C57BL/6JRj* | The Janvier Laboratory | MGI:2670020 |

#### Antibodies

| Reagents | Source | Catalogue, Identifier |
| --- | --- | --- |
| Rabbit phospho-H3 | Cell Signaling Technologies | Cat# 9701, RRID:AB_331535 |
| Anti-rabbit IgG conjugated with AF594 | Thermo Fisher Scientific | Cat#A-11012, RRID:AB_2534079 |
| Rabbit Cleaved Caspase 3 | Cell Signaling Technologies | Cat#9661, RRID: AB_2341188 |
| Rabbit Caspase 3 | Cell Signaling Technologies | Cat#9662, RRID: AB_331439 |
| Mouse Puromycin | Millipore Sigma | Cat#MABE343, RRID:AB_2566826 |
| Rabbit-Hypusinated EIF5A | Millipore Sigma | Cat#ABS1064, RRID:AB_2631138 |
| Mouse EIF5A | BD Biosciences | Cat#611977, RRID:AB_399398 |
| Mouse-S6 ribosomal protein | Cell Signaling Technologies | Cat# 2317, RRID: AB_2238583 |
| Mouse-Ubiquitin P4D1 | Santa Cruz Biotechnology | Cat#sc-8017, RRID: AB_628423 |
| Rabbit-Ubiquitin (linkage specific K48) | Abcam | Cat#ab140601, RRID: AB_2783797 |
| Rabbit-SQSTM1/p62 | Abcam | Cat#ab91526, RRID: AB_2050336 |
| Rabbit-ornithine aminotransferase (ODC1) | Abcam | Cat#ab137679, RRID: AB_2924872 |
| Anti-Rabbit immunoglobulins | Agilent Dako | Cat#P0448, RRID:AB_2617138 |
| Anti-Mouse immunoglobulins | Agilent Dako | Cat#P0447, RRID:AB_2617137 |

#### Chemicals, Peptides, and Recombinant Proteins

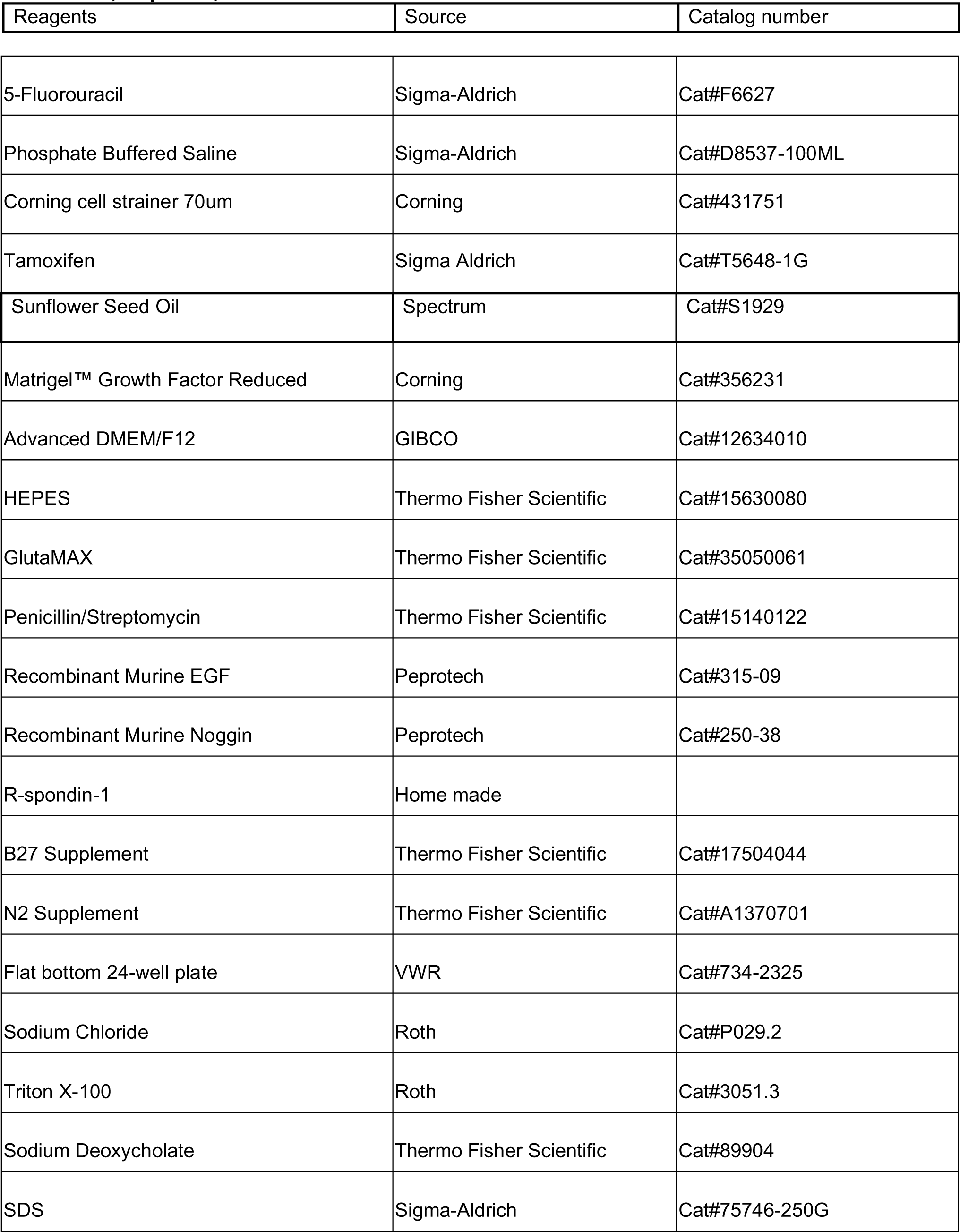
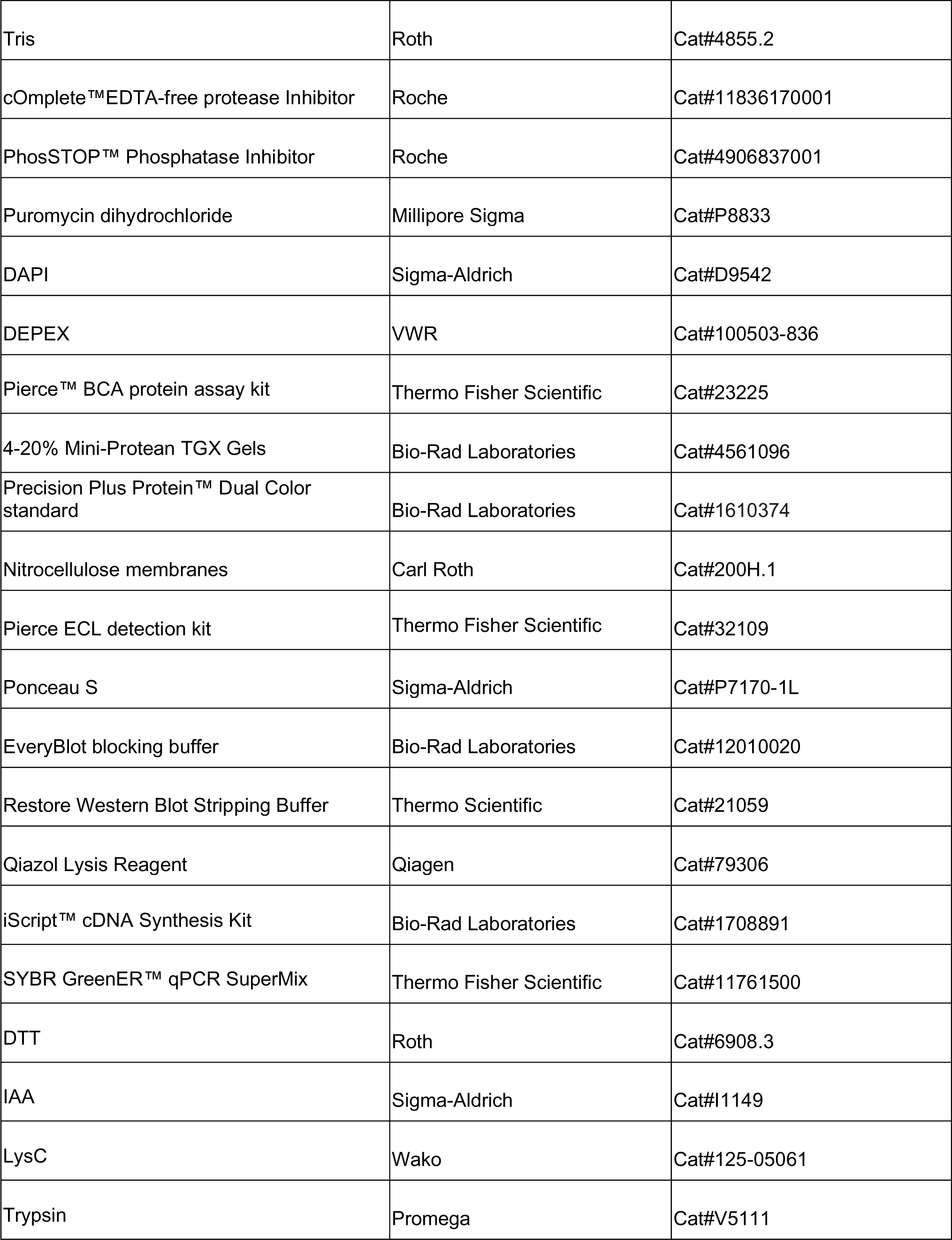
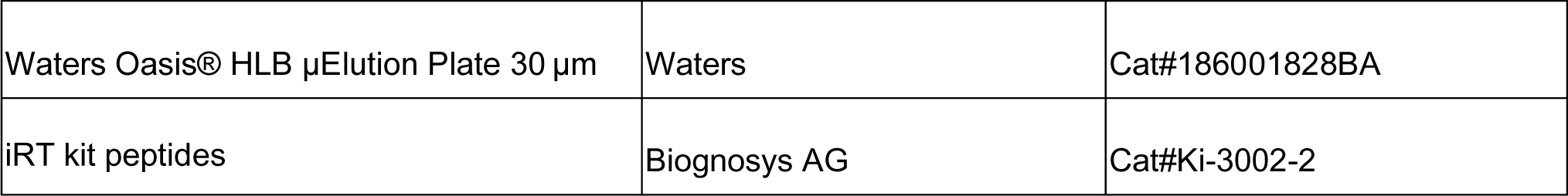

#### Chemicals used for polyamines measurement

| Reagents | Source | Origin |
| --- | --- | --- |
| Acetonitrile | Avantor International | Darmstadt, Germany |
| Acetone | Avantor International | Darmstadt, Germany |
| Methanol | Avantor International | Darmstadt, Germany |
| Acetic acid | Thermo Fisher Scientific | Schwerte, Germany |
| Formic acid | Thermo Fisher Scientific | Schwerte, Germany |
| Ammonium formate | Thermo Fisher Scientific | Schwerte, Germany |
| PBS | CLS | Eppelheim, Germany |
| Dansylchloride | Sigma-Aldrich | Steinheim, Germany |
| Argininosuccinic acid | Sigma-Aldrich | Steinheim, Germany |
| Cadaverin | Sigma-Aldrich | Steinheim, Germany |
| L-anserine | Sigma-Aldrich | Steinheim, Germany |
| L-arginine | Sigma-Aldrich | Steinheim, Germany |
| L-aspartic acid | Sigma-Aldrich | Steinheim, Germany |
| L-citrulline | Sigma-Aldrich | Steinheim, Germany |
| L-ornithine | Sigma-Aldrich | Steinheim, Germany |
| N1,N12-diacetylspermine | Sigma-Aldrich | Steinheim, Germany |
| N1-Ac-putrescine | Sigma-Aldrich | Steinheim, Germany |
| N1-Ac-spermidine | Sigma-Aldrich | Steinheim, Germany |
| N1-Ac-spermine | Sigma-Aldrich | Steinheim, Germany |
| Proline | Sigma-Aldrich | Steinheim, Germany |
| Putrescine | Sigma-Aldrich | Steinheim, Germany |
| Spermidine | Sigma-Aldrich | Steinheim, Germany |
| Spermine | Sigma-Aldrich | Steinheim, Germany |
| Spermine-<sup>2</sup>H<sub>8</sub> | Sigma-Aldrich | Steinheim, Germany |
| L-Arginine<sup>13</sup>C<sub>6</sub> | Cambridge Isotope Laboratories | Tewksbury, MA, USA |
| Spermidine-<sup>2</sup>H<sub>6</sub> | Cayman Chemicals | Hamburg, Germany |

#### Deposited Data

|  |  |  |
| --- | --- | --- |
| Mass spectrometry based proteomics data | MassIVE<br>(https://massive.ucsd.edu) | Proteomic dataset: MSV000094212 |

#### Software and Algorithms

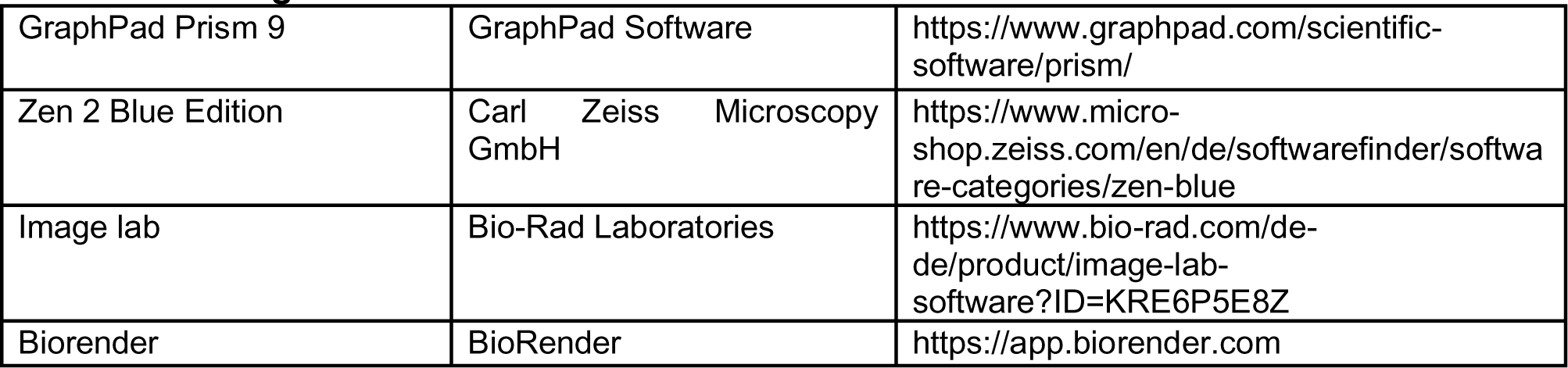

|  |  |  |
| --- | --- | --- |
| GraphPad Prism 9 | GraphPad Software | https://www.graphpad.com/scientific-software/prism/ |
| Zen 2 Blue Edition | Carl Zeiss Microscopy GmbH | https://www.micro-shop.zeiss.com/en/de/softwarefinder/software-categories/zen-blue |
| Image lab | Bio-Rad Laboratories | https://www.bio-rad.com/de-de/product/image-lab-software?ID=KRE6P5E8Z |
| Biorender | BioRender | https://app.biorender.com |

